# Fluorescent peptomer substrates for differential degradation by metalloproteases

**DOI:** 10.1101/2022.08.31.506126

**Authors:** Mariah J. Austin, Hattie C. Schunk, Carolyn M. Watkins, Natalie R. Ling, Jeremy M. Chauvin, Logan D. Morton, Adrianne M. Rosales

**Affiliations:** McKetta Department of Chemical Engineering, University of Texas at Austin, Austin, TX USA; Department of Biomedical Engineering, University of Texas at Austin, Austin, TX USA

## Abstract

Proteases, especially MMPs, are attractive biomarkers given their central role in both physiological and pathological processes. Distinguishing MMP activity with degradable substrates, however, is a difficult task due to overlapping substrate specificity profiles. Here, we developed a system of peptomers (peptide-peptoid hybrids) to probe the impact of non-natural residues on MMP specificity for a MMP peptide consensus sequence. Peptoids are non-natural, *N*-substituted glycines with a large side chain diversity. Given the presence of a hallmark proline residue in the P3 position of MMP consensus sequences, we hypothesized that peptoids may offer *N*-substituted alternatives to generate differential interactions with MMPs. To investigate this hypothesis, peptomer substrates were exposed to five different MMPs, as well as bacterial collagenase, and monitored by fluorescence resonance energy transfer and liquid chromatography-mass spectrometry to determine the rate of cleavage and the composition of degraded fragments, respectively. We found that peptoid residues are well-tolerated in the P3 and P3’ substrate sites and that the identity of the peptoid in these sites displays moderate influence on the rate of cleavage. However, peptoid residues were even better tolerated in the P1 substrate site where activity was more strongly correlated with sidechain identity than sidechain position. All MMPs explored demonstrated similar trends in specificity for the peptomers but exhibited different degrees of variability in proteolytic rate. These kinetic profiles served as “fingerprints” for the proteases and yielded separation by multivariate data analysis. To further demonstrate practical application of this tunability in degradation kinetics, peptomer substrates were tethered into hydrogels and released over distinct timescales. Overall, this work represents a significant step toward the design of probes that maximize differential MMP behavior and presents design rules to tune degradation kinetics with peptoid substitutions, which has promising implications for diagnostic and prognostic applications using array-based sensors.

## Introduction

Matrix metalloproteinases (MMPs) are a family of proteases largely responsible for turnover of extracellular matrix (ECM) as well as other regulatory cellular functions. Their ability to catalyze hydrolysis of various proteins make them key contributors to normal physiological processes like embryogenesis, bone growth, angiogenesis, wound healing, and tissue regeneration,^1,2^ as well as pathological processes including cancer and other inflammatory diseases.^3,4^ Distinguishing MMP behavior is key to understanding the dynamic, cascading pathways that dictate matrix remodeling in healthy and diseased tissue. Moreover, there is considerable interest in the development of selective substrates to improve the function of MMPs as biomarkers for diagnostic and prognostic assays,^5–7^ tissue engineering scaffolds,^8–13^ therapeutic linkers,^14–16^ and molecular inhibitors.^17,18^ Despite this large body of work, design rules for developing targeted degradable peptide sequences remain uncertain.

Current approaches to distinguish MMP activity suffer from overlapping cleavage specificity profiles.^19,20^ Numerous proteomic studies have taken a comprehensive approach toward analyzing the factors that influence enzyme-substrate specificity with the goal of identifying highly specific substrates.^21–23^ Many utilize proteomic identification of cleavage sites (PICS) wherein specificity profiles are deduced from the cleavage products of native protein fragments, and consensus sequences are then established according to the most frequently observed amino acids in each substrate position.^24–26^ These consensus sequences, however, are not optimized to uniquely target individual enzymes and are therefore susceptible to off-target degradation and limit the accuracy of peptides as biosensing probes.^7^

Given this limitation, some studies have explored substrates with non-natural residues to leverage alternative backbone interactions and side chains that may generate substrates uniquely recognized by individual proteases.^27–29^ Drag and colleagues incorporated non-natural amino acids into combinatorial libraries to profile the specificity of caspases,^30^ cathepsin G,^31^ and cathepsin L.^32^ These studies demonstrated that a non-natural substitution in a single substrate site can enhance selectivity for certain proteases in the same family. Extending beyond the monomer structure of amino acids, *N*-alkylation of a single residue was proven to be effective in increasing substrate resistance to elastase.^33^ While there was still overlap in specificity for single enzyme-substrate pairs, this prior work suggests that non-natural residues are a promising strategy to generate variance in proteolytic susceptibility.

Here, in a complementary approach, we aimed to strategically alter the affinity of MMP substrate sequences using *N*-substituted peptoids and leverage cross-reactivity to elicit distinct patterns in degradation between similar proteases. We were especially intrigued by the hallmark proline residue consistently found in the P3 position of MMP degradable substrates. Proline is the only natural amino acid with a tertiary amide, a feature shared by peptoids. Peptoids are *N*-substituted glycines, where the sidechain is appended to the nitrogen on the polyamide backbone. This shift of attachment point by one atom relative to peptides eliminates the chirality at the α-carbon and removes the secondary amide hydrogen responsible for backbone hydrogen bonding that stabilizes higher order structure and intermolecular complexation. Therefore, peptoids introduce conformational flexibility,^34^ and they have been shown to be resistant to degradation by common proteases.^35,36^ Peptoids provide straightforward means for probing differences in MMP cleavage behavior because they can be synthesized with the same sidechains as natural amino acids, thereby preserving the overall chemical nature of the substrate and allowing for systematic study of side chain versus backbone hydrogen bonding interactions during enzymatic cleavage.^37^ In addition, their submonomer synthesis^38^ can be easily integrated with traditional solid phase peptide synthesis to produce peptide-peptoid hybrids, termed “peptomers.”^39^

Peptomers with peptoids in the P1’ position have previously been designed as inhibitors for numerous proteases of various types and origins.^40–43^ Beyond the P1’ position, however, it is unclear how peptoids influence enzyme recognition and catalytic efficiency. In one report, Stawikowski *et al*. incorporated peptoid residues in the P3, P1, P1’, and P10’ positions of collagen triple-helical peptides and examined their susceptibility to MMP-1, MMP-8, MMP-13, and MT1-MMP.^44^ Interestingly, their study identified a sequence with high MMP-13 selectivity where the scissile site shifted due to a unique interaction between MMP-13’s hemopexin domain and the peptoid residue in the resulting P7’ position. These results confirm that peptoid residues influence substrate recognition by MMPs and suggests the proteolytic resistance of *N*-substitutions can be harnessed to generate differential degradation behavior.

In this study, we first investigated the impact of individual peptoid substitutions on substrate degradability, then engineered selected peptomer substrates for cross-reactive sensing formats by creating “fingerprints” with high discriminatory power for MMPs. Adapting well-established methods for protease substrate profiling,^45–48^ we synthesized methoxycoumarin and dinitropheynyl-labeled substrates to monitor their cleavage via fluorescence resonance energy transfer, as well as by liquid chromatography and mass spectrometry. We started with a PICS derived pan-MMP consensus sequence, PAN↑LVA (henceforth referred to as the *Pan-MMP Peptide*),^26^ to readily compare changes in proteolysis among similar proteases: Collagenase Type I from *Clostridium histolyticum* and recombinant human MMPs, namely the collagenases (MMP-1, MMP-8, MMP-13), and gelatinases (MMP-2, MMP-9). Using this parent sequence, we took a systematic approach to understand how peptoid substitutions impact cleavage behavior, using two small libraries of substrates with peptoid substitutions adjacent to the scissile site: 1) a peptoid analog library with translocated sidechains that match the amino acids they were replacing in each substrate position, and 2) a similarity scan based on sarcosine substitutions. These libraries yielded crucial information about which residues were critical to proteolysis, which we used to develop substrates with multiple strategically placed peptoid substitutions to further understand and direct MMP cleavage behavior. We characterized these substrates using circular dichroism and Michaelis-Menten parameter approximations to elucidate the molecular underpinnings of the proteolytic behavior observed. Excitingly, this array of substrates exhibited differential cleavage rates, which were directly processed by multivariate data analysis techniques and implemented in a pattern-recognition algorithm to demonstrate how peptomers can be used to distinguish between MMPs. In addition, a subset of the tunable peptomers were incorporated as fluorescent reporters in a hydrogel network, demonstrating the customizable degradation rate enabled with peptoid residues. Altogether, our findings establish peptomers as tools for discriminating between proteases with similar specificity by relying on the composite response of all substrates in the array, as well as expand the available strategies for controlling degradation in biomaterial platforms.

## Experimental

### Materials

L-amino acids, sarcosine, and modified lysines were all Fmoc-protected and purchased from Chem-Impex International, Inc. (Wood Dale, IL), along with Rink amide resin, *O*-(1H-6-Chlorobenzotriazol-1-yl)-*N,N,N*’,*N*’-tetramethyluronium hexafluorophosphate (HCTU, ≥ 99%), bromoacetic acid (≥ 99%), *N,N’*-Diisopropylcarbodiimide (DIC, ≥ 99%), and *N,N’*-Diisopropylethylamine (DIPEA, 99.8%). Peptoid submonomers, including 1-(2-aminoethyl) pyrrolidine (98%), isopropylamine (99.5%), isobutylamine (99%), and glycinamide hydrochloride (98%) were purchased from Acros Organics (Fir Lawn, NJ), as well as triisopropylsilane (TIPS, 98%). 2-phenethylamine (99%) was from Alfa Aesar (Haverhill, MA). *N*-methylmorpholine (NMM, 99%), piperidine (99.5%), and 4-Aminophenylmercuric acetate (APMA, ≥97%) were purchased from Millipore Sigma (Burlington, MA). Sodium hydroxide (NaOH, ≥97%) and all solvents were purchased from Fisher Scientific (Hampton, NH) at the following purity levels: dimethylformamide (DMF, ≥99.8%), trifluoroacetic acid (TFA, ≥99.5%), diethyl ether (ether, ≥99%), acetonitrile (ACN, ≥99.9%), 2-propanol (IPA, ≥99.5%), dimethylsulfoxide (DMSO, ≥99.7%).

Full-length recombinant, carrier-free human matrix metalloproteinases were purchased from R&D Systems, Inc. (Minneapolis, MN) (**Table S1**). Gibco® Collagenase, Type 1, from *Clostridium histolyticum* was purchased as a lyophilized powder from Thermo-Fisher Scientific (Waltham, MA) with a specific activity of 250 units mg^-1^. MMP inhibitor, GM 6001 (>95%), was manufactured by Calbiochem and purchased from Millipore Sigma. Tris-HCl (≥ 99%) and sodium chloride (NaCl, ≥ 99%) were purchased from Fisher Scientific. Brij®-35 and calcium chloride (CaCl_2_, ≥ 96%) were purchased from Acros Organics.

4-arm 10kDa poly(ethylene glycol)-amine (PEG-amine, ≥ 95%) was purchased from JenKem Technology USA (Plano, TX). 5-norbornene-2-carboxylic acid (98%) and 3.4 kDa linear PEG dithiol were purchased from Millipore Sigma. Lithium phenyl(2,4,6-trimethylbenzoyl)phosphinate (LAP, 98%) and *O*-(Benzotriazol-1-yl)-*N,N,N*’,*N*’-tetramethyluronium hexafluorophosphate (HBTU, ≥ 98%) were manufactured by TCI America and purchased from Fisher Scientific.

### Methods

#### Substrate synthesis

All reagents were used as purchased, with the exception of glycinamide HCl, which was free-based using NaOH in IPA according to a previously reported protocol.^49^ Peptides, peptoids, and peptomers were all synthesized using Rink Amide polystyrene resin (0.43 mmol g^-1^) on a Prelude X automated peptide synthesizer (Gyros Protein Technologies) at a scale of 50 μmol. Fmoc groups were removed from the resin and subsequent amino acids by washing twice with 20% piperidine in DMF. Peptide residues and sarcosine utilized Fmoc-protected amino acids (250 mM, 5x molar excess) coupled using HCTU activator (250 mM, 5x molar excess) and NMM (500 mM, 10x molar excess). Coupling steps were performed twice. Peptoid residues were installed according to previously published submonomer synthesis methods.^38,50^ First, bromoacylation occurs via addition of bromoacetic acid (1.2 M in DMF) and DIC at a molar ratio of 1:0.93. The bromine is then displaced by a primary amine (2M in NMP) to install the entire peptoid residue. Upon completion of synthesis, substrates were cleaved from the resin using a cleavage cocktail comprised of 95% TFA/ 2.5% water/ 2.5% TIPS for substrates with a sidechain protecting group (those with Asn(Trt)), or 95% TFA/ 5% water. The resin was then filtered off, and the substrates were prepared for purification.

#### Substrate purification

Crude purification by ether precipitation was possible for all substrates except the *Peptoid*. Substrates dissolved in cleavage cocktail were added dropwise to a ten-fold volume excess of chilled ether, centrifuged to collect the precipitate, and then washed twice with fresh ether. For the *Peptoid*, the cleavage cocktail was evaporated on a rotary evaporator. Substrates were dried in a vacuum oven overnight to remove trace ether, then dissolved in a mixture of 25% acetonitrile/ 75% water with 0.1% TFA and purified with a semi-prep C18 column on a Dionex UltiMate 3000 UHPLC with a 15 minute gradient from 25-100% acetonitrile at 10 mL min^-1^. Fractions were collected by their UV signal at 214 nm and the samples were re-purified until only a single peak remained (1-3 purification cycles). Purified substrates were then lyophilized and analyzed by HPLC with an analytical C18 column and via matrix-assisted laser desorption/ionization-time of flight (MALDI-TOF) mass spectrometer using a Bruker autoflex maX instrument to assess purity and confirm molecular weight, respectively (**Figure S1**).

#### Fluorescence assays

Substrate cleavage was monitored in real-time by tracking the fluorescent signal of 7-methoxycoumarin-4-acetic acid as it was liberated from the dinitrophenyl quencher by enzymatic hydrolysis. Substrates were first dissolved in DMSO at a concentration of 1 mM, as measured by the dinitrophenyl group’s absorbance at 363 nm on a NanoDrop OneC Microvolume UV-Vis Spectrophotometer using an experimentally derived extinction coefficient of ε_363 nm_= 16,900 cm^-1^ M^-1^. Substrates were then diluted to 20 μM with 10% DMSO in MMP buffer (50 mM Tris HCl, 10 mM CaCl_2_, 150 mM NaCl, 0.05% (w/v) Brij®-35, pH 7.5). MMPs were activated according to procedures provided by the supplier using APMA, then diluted to the concentration of interest in buffer. Next, 50 μL of substrate solution was combined with either 50 μL of buffer (controls) or 50 μL of enzyme solution (samples) in triplicate in a black 96-well plate. The plate was oscillated for 10 seconds to mix and then read on a BioTek Synergy H1 Multi-Mode Microplate Reader at Ex./Em. 325/392 nm for three hours. Fluorescence values of the controls were subtracted from the sample wells for fluorescence plots.

#### Statistical analysis of cleavage rates

Cleavage rates were compared by fitting a simple linear regression to the first ten time points of fluorescence traces using GraphPad Prism 9. One-way Analysis of Variance (ANOVA) was conducted to individually determine the degree of significant difference between cleavage rates of different substrates. When measuring rate changes, two-way ANOVA was applied to determine if the change was statistically different than zero.

#### Degradation detection

Fractions of cleaved substrates were measured after 24 hours using liquid chromatography-mass spectrometry (LC-MS). 10 μM of substrate in 5% DMSO was incubated for 24 hours at 37°C with the same enzyme concentrations used for fluorescence assays. Controls without enzyme were prepared in the same way. After 24 hours, the reactions were quenched in liquid nitrogen, then lyophilized. Lyophilized samples were reconstituted at 50 μM in water with 25% acetonitrile and analyzed on an Agilent Technologies 6120 Single Quadropole LC-MS with an Agilent ZORBAX Eclipse Plus C18 column running acetonitrile and water with 0.1% formic acid. A 12 minute gradient ramp spanned from 5% to 100% acetonitrile. Spectra were extracted in Agilent’s ChemStation software at 210 nm and 400 nm, along with extracted ion chromatographs at each relevant fragment mass in positive ion mode. The fragments in the 400 nm trace (only those with a dinitrophenyl group) were matched to their extracted mass peak at the same retention time and integrated to determine the composition of fragments after cleavage. Only one sample was analyzed using LC-MS; however, fluorescence traces serve as a secondary means of validating the data.

#### Circular dichroism (CD)

CD spectra were acquired from 190 to 260 nm on a Jasco J-815 CD Spectropolarimeter at 25 °C. Samples were dissolved in 30% acetonitrile in water at approximately 500 μM. Samples were diluted until the high tension (HT) voltage was less than 600 volts at the lowest wavelength to prevent signal saturation. After measurement, samples were measured on NanoDrop OneC Microvolume UV-Vis Spectrophotometer to accurately quantify concentration. Mean residual ellipticity was then calculated according to the following equation:

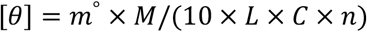

where *m°* is the CD given in millidegrees, *M* is the molecular weight (g mol^-1^), *L* is the path length of the cell (cm), *C* is the concentration of substrate (g L^-1^), and *n* is he number of monomers.

#### Active site titrations

MMPs were active site titrated by inhibition assay using tight-binding inhibitor, GM 6001. First, activated MMPs were incubated with inhibitor (0-20 nM) for 20 minutes at 37°C. Next, substrate solution was added to a final concentration and volume of 10 μM and 100 μL, respectively. Substrate cleavage was fluorescently monitored for 15 minutes. Initial velocities were determined by fitting the traces to a simple linear regression using GraphPad Prism 9. The initial velocities were then plotted against inhibitor concentration and modeled with the Morrison equation modified for fractional rates:^51–53^

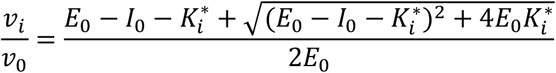

where *v*_*i*_ is the initial reaction velocity at a certain inhibitor concentration, *v*_*0*_ is the uninhibited initial reaction velocity, *E*_*0*_ is the concentration of enzyme active sites, *I*_*0*_ is the concentration of inhibitor, and 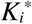 is the applied dissociation constant. *E*_*0*_ and 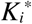 were fit parameters. Fitting was accomplished in GraphPad Prism 9, which eliminated outliers for parameter fitting using a robust nonlinear regression Q coefficient of 1%.

#### Michaelis-Menten kinetic parameter determination

Kinetic parameters were estimated by two methods: 1) using a publicly available R package developed by Choi *et al*. that models enzyme kinetics using the Metropolis-Hasting algorithm with Gibbs sampler based on Bayesian inference,^54^ and 2) a low substrate-concentration approximation based on first-order reaction kinetics. To do so, a subset of key substrates were fluorescently screened at 0.5 μM for three hours. Progress curves were fit to the following equation:

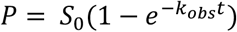

where *P* is the concentration of product (cleaved substrate), *S*_*0*_ is initial substrate concentration, and *k*_*obs*_ is the exponential fitting parameter.^55^ At low substrate concentrations ([S]<<K_M_), when the Michaelis-Menten equation is in the first-order reaction phase *k*_*obs*_ is proportional to the catalytic efficiency as follows:

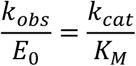

where *E*_*0*_ is enzyme concentration (determined by active site titration), *k*_*cat*_ is the turnover number, *K*_*M*_ is the Michaelis constant, and the ratio of *k*_*cat*_ to *K*_*M*_ is catalytic efficiency.^56^ Progress curves at 10 μM and 0.5 μM were both used for parameter estimations in the R code.

#### Preprocessing and multivariate data analysis

10 μM fluorescence traces were fit to the following built-in exponential plateau function in GraphPad Prism 9:

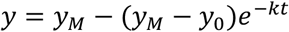

The *y* inputs remained in their raw data form of relative fluorescence units (RFU). Other parameters are defined as follows: *y*_*M*_ is the plateau value, which was constrained to a value of 530,000 RFU for all fits (corresponding to complete hydrolysis and maximum fluorescence signal), *y*_*0*_ is the initial value, which was constrained to zero for all fits, and *k* (s^-1^) represents the rate of reaction. The logarithm of the k-constants were used as features for subsequent multivariate data analysis.

The data matrix was processed using both unsupervised and supervised techniques. For hierarchal clustering analysis (HCA), the MATLAB function ‘linkage’ was used with ‘cityblock’ distance metric and ‘average’ linkage metrics. The function ‘dendrogram’ was used for visualization. For principal component analysis (PCA), the data matrix was directly fed to the ‘pca’ function in MATLAB, and the outputs ‘coeff,’ ‘score,’ ‘latent,’ and ‘explained’ were used to construct visuals in lower dimensional space.

Following unsupervised analysis, the first two principal components were used to train and test the classification accuracy using the k-nearest neighbor (kNN) model. The MATLAB ‘fitcknn’ and ‘kfoldpredict’ functions were used with k=2 neighbors and three-fold, stratified cross-validation to evaluate the classification accuracy.

#### PEG-Norbornene functionalization

4-arm PEG-norbornene was synthesized by activating 5-norbornene-2-carboxylic acid (0.244 mL, 2 mmol, 5.0 equiv) with HBTU/HOBt (834.4 mg, 2.2 mmol, 5.5 equiv) for 3 minutes in DMF (5 mL) under an argon purge. DIPEA (0.418 mL, 6.0 equiv) was added to this activated solution and stirred for 5 minutes at room temperature. Next, this solution was added to an argon purged flask containing 4-arm, 10 kDa PEG-amine (1.0 g, 1.0 equiv), and the reaction proceeded under stirring for 24 hours at room temperature. Upon completion of the reaction, the product was precipitated in cold ether (60 mL) and subsequently washed twice with fresh ether, then dialyzed for 3 days in distilled water. The product was recovered as a white solid. Yield = 93%, functionalization = 91.5% (**Figure S2**). ^1^H NMR (D_2_O, 400 MHz): δ 6.20 to 5.86 (m, 2H), δ 3.65 to 3.40 (m, 227H).

#### Hydrogel formulation

PEG gels were formed by combining a 4 wt.% solution of PEG-norbornene with PEG-dithiol at a thiol:ene ratio of 0.95:1 and cysteine-tagged fluorescent substrate at a thiol:ene ratio of 0.05:1. Thus, the total weight percent of PEG was 6.3% in water with 10% DMSO to prevent substrate precipitation from solution. LAP photoinitiator was included at 0.05 wt.% and gelation was induced by exposure to 365 nm light (10 mW cm^-2^, 160 seconds). Gels were 30 μL formed in 24-well plates. Blank gels were formed following the same procedure with PEG-dithiol at a thiol:ene ratio of 0.95:1, but without cysteine-tagged fluorescent substrates.

#### Hydrogel release experiments

Gels were swollen in 0.5 mL of buffer with 10% DMSO twice for 24 hours at a time to remove any untethered substrate and equilibrate the gels. After the two swell/wash steps, gels were treated with either:1) MMP buffer with 10% DMSO for controls and blanks, or 2) MMP buffer with 10% DMSO and MMP for samples. MMP-2 was added at 0.13 μg mL^-1^, and MMP-9 was added at 0.5 μg mL^-1^. Fluorescence measurements were taken at predetermined time intervals between 0 and 72 hours using an area scan to track cleavage of fluorescent substrates by MMPs and their subsequent diffusion out of the gel. Fluorescence measurements were taken on a BioTek Synergy H1 Multi-Mode Microplate Reader at Ex./Em. 325/392 nm. An area scan was performed using a 24 well plate format with a 7×7 matrix. All experimental conditions were performed in triplicate.

#### Controlled-release data processing

The region of interest (ROI) for each area scan was the three squares with highest intensity in the 7×7 matrix, corresponding to the center of the gel where substrate was tethered. This ROI for each gel was determined at t=0 and maintained for all subsequent analysis. The fluorescence intensity in the ROI of the blank gels was averaged at each timepoint and subtracted from the control and MMP gels to account for background fluorescence. Next, each sample was normalized by the fluorescence intensity of the t=0 timepoint, representing the fraction of substrate remaining in the gel. The replicates were then averaged and finally, the averaged MMP-2 and MMP-9 samples were divided by the averaged controls at each timepoint to account for fluorescence fluctuations and any substrate diffusion not due to proteolysis. The trace for each enzyme-substrate combination was fit to GraphPad Prism 9’s built-in nonlinear regression one-phase decay model (see above for variable descriptions):

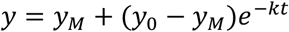

Half-lives (hours) were calculated as *ln(2)/k* and are reported with their 95% confidence interval.

## Results and Discussion

### Peptoid analogs to explore the role of N-substitution by substrate position

To establish a baseline for comparison, we first assessed the cleavage of the *Pan-MMP Peptide* sequence, PAN↑LVA (**Figure 1A**), by bacterial collagenase, MMP-1, MMP-8, MMP-13, MMP-2, and MMP-9. We included bacterial crude collagenase because it is often used as an inexpensive substitute to model MMP behavior in biomaterial studies.^57,58^ We hypothesized that the crude collagenase would demonstrate similar trends as the MMPs with less specificity because it could contain a heterogeneous mixture of proteases with varying activities.^59^ For the selected set of MMPs, they represent key proteases that are often dysregulated in pathological processes, making them attractive biomarkers.^3^ The substrate was exposed to each of the enzymes at equivalent mass concentrations (0.5 μg mL^-1^), and fluorescently monitored to determine the rate of cleavage (**Figure 1B**). Unsurprisingly, the enzymes demonstrated varied activity with MMP-13 and MMP-2 cleaving the fastest. In order to compare the peptomer substrates’ responses to the original peptide sequence, we normalized the concentration of each MMP to cleave the *Pan-MMP Peptide* at the same rate, ensuring it reached saturation within the three-hour time course of our experiment (**Figure 1C**). All subsequent experiments were conducted using these activity-normalized concentrations, although to compare relative catalytic efficiency, we later determined the Michaelis-Menten parameters.

**Figure 1:**
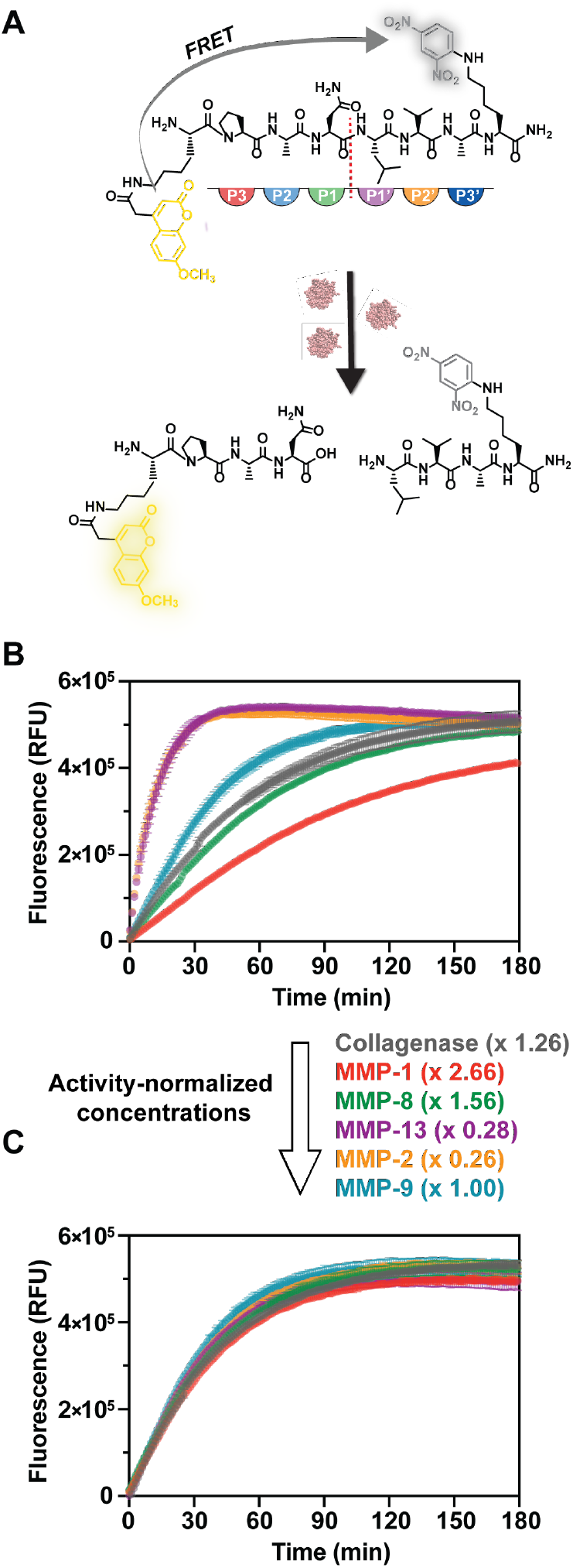
*Pan-MMP Peptide* cleavage rates by collagenase and MMPs. (A) *Pan-MMP Peptide* consensus sequence with fluorophore and quencher moieties attached (top). Enzyme-substrate interactions are described according to the protein/peptide position (P) which match corresponding pockets in the enzyme binding site. Beginning at the scissile site, residues increase in number toward the substrate termini and are termed as either: 1) the prime motif toward the C-terminus, or 2) non-prime toward the N-terminus. When the substrate is intact, the fluorophore is quenched. Upon cleavage, the fluorophore is liberated from the quencher and produces a quantitative fluorescent output. (B) Fluorescence tracking of *Pan-MMP Peptide* degraded by MMPs at the same concentration (0.5 μg mL^-1^) and (C) at activity-normalized enzyme concentrations. Error bars represent standard deviation from three technical replicates.

We next investigated the impact of *N*-substitutions at each residue position on MMP degradation rate. We began with a peptoid analog library with translocated side chains that matched the *Pan-MMP Peptide* amino acid sequence. The peptoid analog library included six individually substituted peptomers (named by the position and amino acid being translocated:^60^ *P3 NPro, P2 NAla, P1 NAsn, P1’ NLeu, P2’ NVal*, and *P3’ NAla*) and a fully substituted peptoid (*Peptoid)* with *N*-substituted residues in each substrate active site position (**Figure 2A**). Each substrate was eight residues in length, six of which were the active sequence, with fluorophore and quencher moieties attached as modified lysines on each end.^45^ The peptoid substitutions made at each active sequence position were synthesized with sidechains identical to those of the amino acids they replaced, except in the case of the P3 position. In that position, we used N-[2-(1-Pyrrolidinyl)ethyl] glycine (*NPro*) which consists of the same pyrrolidine ring as the native proline residue without the bond to the backbone α-carbon.

**Figure 2:**
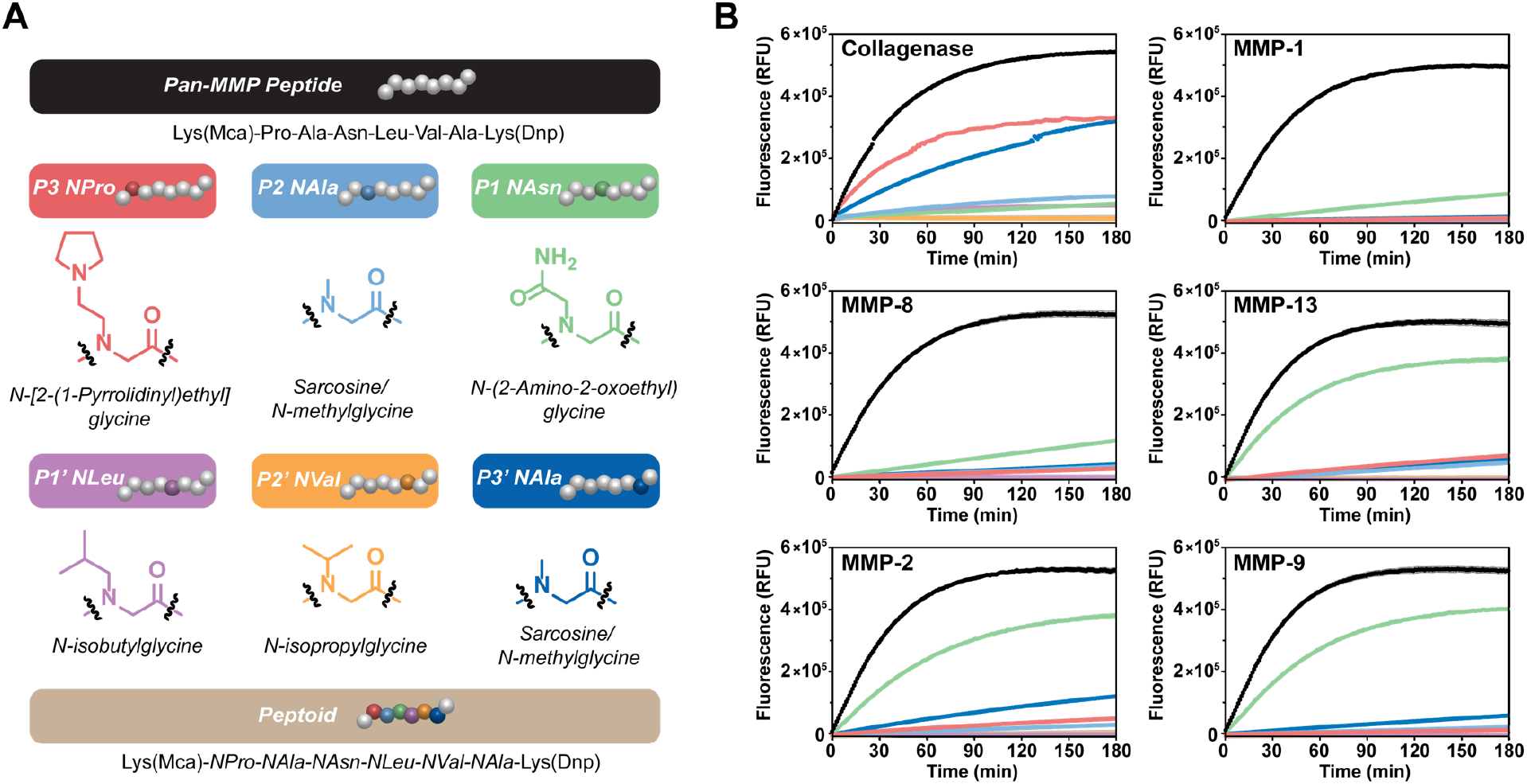
Peptoid analog library. (A) Original *Pan-MMP Peptide* sequence with monomer structures of the *N*-substitutions used to make peptomers and a full peptoid, along with cartoon depictions of the substrates where colored bubbles represent a peptoid residue. (B) Fluorescence traces measuring substrate cleavage by collagenase from *Clostridium histolyticum* and MMPs. Error bars represent standard error of the mean from three technical replicates.

The peptoid analog substrates were exposed to collagenase and the five MMPs and tracked fluorescently (**Figure 2B**). The bacterial collagenase was able to cleave all the substrates except the *P2’ NVal* and *Peptoid*. Similarly, none of the MMPs cleaved the *P1’ NLeu, P2’ NVal*, or *Peptoid* substrates to a measurable degree. However, the substrate order of activity varied between the collagenase and MMPs. Collagenase cleaved *P3 Pro* and *P3’ NAla* fastest, followed by *P2 NAla* and *P1 NAsn*. The MMPs were able to cleave all these substrates, but at rates at least 10-fold slower than the *Pan-MMP Peptide*, with the exception of the *P1 NAsn* peptomer. *P1 NAsn* was cleaved significantly faster than the other peptomers, albeit spanning a range from 6% (MMP-1) to 55% (MMP-13) of the benchmark initial rate measured for the *Pan-MMP Peptide*. Notably, the structural distinction between the gelatinase and collagenase MMPs is that the gelatinase MMPs contain a fibronectin-type domain that enables binding to gelatin.^61,62^ This feature does not appear to play a role in peptomer recognition as MMP-13 behaved very similarly to MMP-2 and MMP-9 here, but it may be relevant for larger substrates that bind beyond the active site. Within the active site, ^61^ the catalytic mechanism proposed for MMPs (**Figure S3**)^63,64^ shows the P1 residue carbonyl co-ordinating with the catalytic zinc ion, but the remainder of the residue does not directly interact with the enzyme active site, suggesting the amide hydrogen in P1 is not critical to binding or hydrolysis. Conversely, the P1’ and P2’ residue amide hydrogens form hydrogen bonds with the enzyme active site, which may be essential for binding. Thus, peptoids substituted into these positions were expected to inhibit or significantly hinder cleavage, explaining why the *P1’ NLeu* and *P2’ NVal* substrates remained fully intact after exposure to the MMPs. To further examine how bacterial collagenase was able to retain activity for these substrates, we investigated the substrate degradation products by LC-MS (**Figure S4**).

The composition of degraded fragments was determined after 24 hours of enzyme exposure at the same activity-normalized concentrations. We compared collagenase (**Figure 3A**) to the highest activity MMP, MMP-2 (**Figure 3B**), and found a key difference in the degradation profiles of the proteases. MMP-2 only cleaved at the predicted cleavage site between the asparagine and leucine residues, whereas collagenase was able to cleave in that position (primary site), as well as the site one position toward the N-terminus (secondary site). A second cleavage site effectively shifts the binding interaction with the substrate, meaning the peptoid residue appears in different positions depending on the scissile site (**Figure S4**). The distribution of cleavage products (*i*.*e*., proportion of primary versus secondary cleavage fragments) was influenced by the location of the peptoid substitution in the chain. For example, bacterial collagenase cleaved the *P1’ NLeu* substrate only at the secondary site, where the peptoid was effectively in the P1 position. Similarly, the *P3’ NAla* substrate only cleaved at the primary cleavage site, where the peptoid was in the P3’ position, and did not cleave at the secondary site, where the peptoid residue was effectively in the unpreferred P2’ location. When comparing the location of peptoid residues in reference to the observed cleavage sites for both enzymes (**Figure 3C**), collagenase and MMP-2 demonstrate similar profiles with the highest fraction of cleavage products when peptoids are in the P3, P1, and P3’ positions. Fluorescence degradation tracking is only sensitive to an initial cleavage event, meaning the rates shown may not reflect the actual proteolytic activity in the case of multiple cleavage sites. Therefore, we discontinued use of the bacterial collagenase for subsequent investigations. In summary, the peptoid analog library suggests that the P1’ and P2’ substrate positions require hydrogen-bonding nitrogens for metalloprotease cleavage and that MMPs are markedly active to peptomers with peptoids in the P1 substrate site.

**Figure 3:**
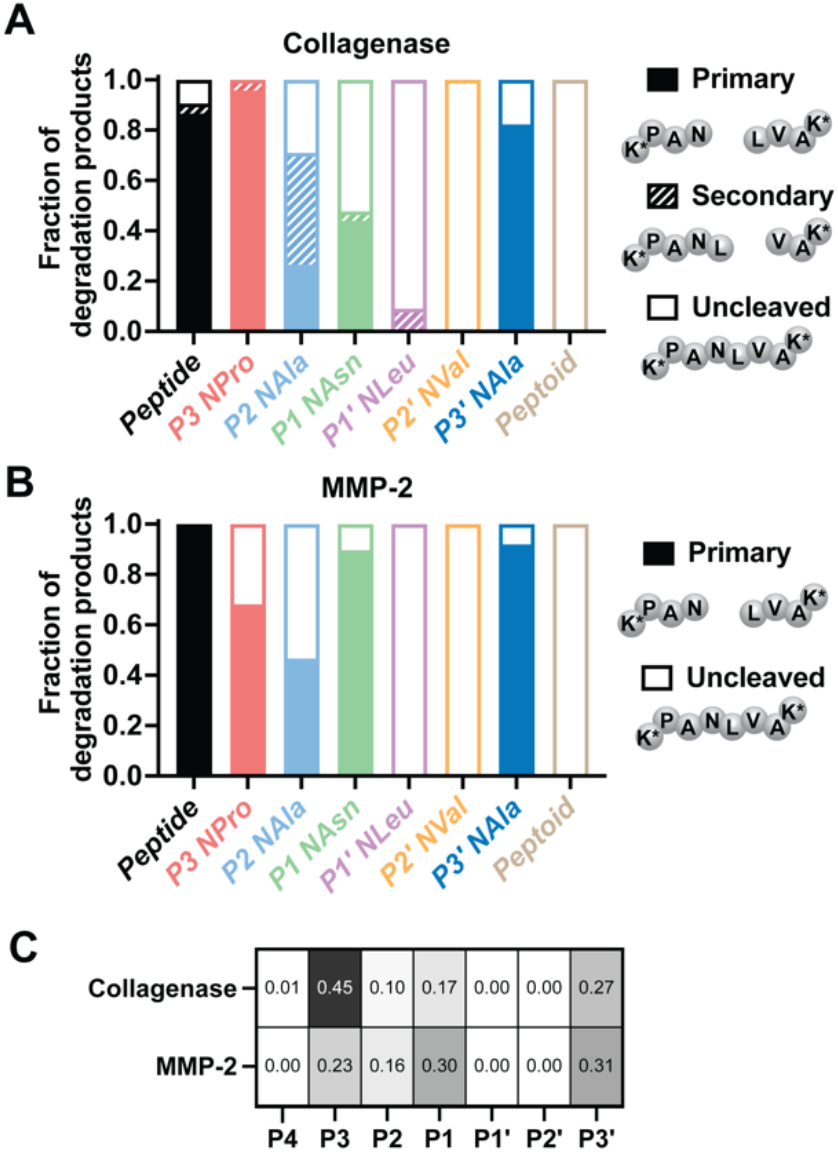
Cleavage site comparison. (A) Hydrolysis fragments measured by LC-MS after 24 hours of incubation with collagenase and (B) MMP-2. (C) Fraction of peptomer cleavage products for each enzyme by peptoid substitution location. LC-MS traces with fragments labeled are shown in **Figure S4**.

### Impact of sidechain identity on peptomer recognition

Upon completion of an initial scan using peptoid analogs, we sought to discern substrate positions where peptoid sidechain identity influences susceptibility to cleavage versus those that inhibit cleavage regardless of identity. To do so, we performed a similarity scan wherein sarcosine was used as the peptoid residue in each substrate position that did not already have it as a peptoid analog (**Figure 4A**). Sarcosine is the smallest peptoid residue and therefore we hypothesized it would minimize the effect of sidechain chemistry on enzyme recognition. In the P3 and P1 positions, using sarcosine increased the cleavage rate for all of the MMPs tested (**Figure 4B**). To quantify the activity difference between the peptoid analogs and the sarcosine scans, the initial rate from each kinetic trace was normalized as a percentage of the benchmark rate of the *Pan-MMP Peptide*. For example, for MMP-1, *P3 NAla* increased the initial rate of cleavage by 15% over the *P3 NPro* substitution (where the *P3 NPro* initial rate was only ∼3% of the *Pan-MMP Peptide*, and *P3 NAla* was 18%). Similarly, *P1 NAla* increased the initial rate for MMP-1 by 20% compared to the *P1 NAsn* substrate. In general, sarcosine significantly increased the initial rate of cleavage in the P3 and P1 positions for all MMPs, with an average increase of 21% for the P3 position and 35% for the P1 position. As expected, *NAla* substitutions in the P1’ and P2’ positions still rendered the substrates resistant to proteolysis.

**Figure 4:**
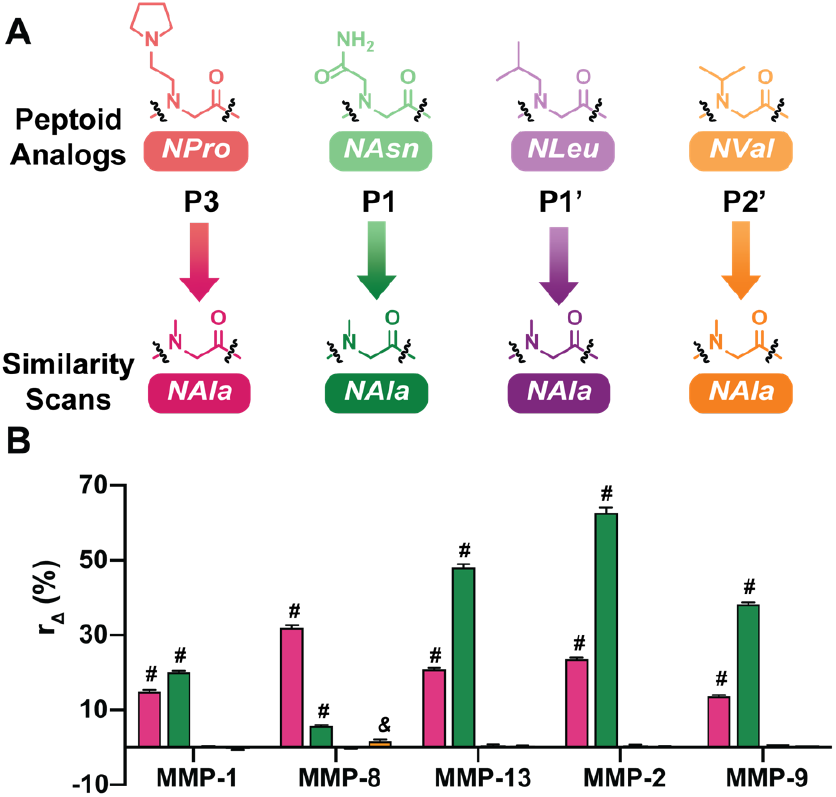
Similarity scan using sarcosine. (A) Substrate positions explored in the similarity scan. Comparisons are between the peptoid analog residue in each position and sarcosine. (B) Change in initial rate of hydrolysis between peptoid analog residue and sarcosine as a percentage of the cleavage rate of the *Pan-MMP Peptide* for each MMP tested. *r*_Δ_ = ((*r*_*sar*_ − *r*_*ana*_)/*r*_*cont*_) × 100 where *r*_*Δ*_ is the rate change, *r*_*sar*_ is the initial rate of the sarcosine scan substrate, *r*_*ana*_ is the initial rate of the peptoid analog substrate and *r*_*cont*_ is the benchmark initial rate of the control peptide. Error bars represent the standard error of the mean. Symbols represent statistical difference from a change of 0%: # = p<0.0001, & = p<0.01. Fluorescence spectra are included in **Figure S4**.

Given these results, we further probed the role of identity versus residue type in the versatile P3 and P1 positions. The hallmark proline residue in MMP consensus sequences made the P3 position especially intriguing for peptoid substitution. In addition to the original *P3 NPro* peptomer and the *P3 NAla* substrate from the sarcosine scan, N-(2-phenylethyl)glycine was used as the monomer to generate the *P3 Npea* substrate. When compared to the *NPro* substitution, *Npea* is similar in size, but significantly more hydrophobic, affording a comparison of the residues’ chemical nature versus sterics (**Figure S5A)**. Similarly, proline is a hydrophilic amino acid while sarcosine is moderately hydrophobic. All of the MMPs preferred the P3 residues in the same order: *Pro* > *NAla* > *NPro* > *Npea*. (**Figure S5B*-*C**). The larger sidechains severely slowed proteolysis, suggesting the size of the residue plays a key role in modulating proteolysis at this position.

We were also intrigued by the especially high tolerance of peptoid substitutions in the P1 site. The *P1 NAla* substrate cleaved faster than the *P1 NAsn* substrate for all of the MMPs. For MMP-13, MMP-2, and MMP-9, the *P1 NAla* substrate cleaved at rates similar to the *Pan-MMP Peptide*, which supports the hypothesis that sidechains in the P1 position do not form any intramolecular contacts with the enzyme and instead may only influence enzyme binding by the steric resistance of fitting into the S1 active site pocket. For the P1 site, we noted that Eckhard *et al*. identified a consensus peptide sequence (PAA↑LVA) for the gelatinase MMPs that resembled our sarcosine substituted sequence.^26^ Thus, we also synthesized this second peptide control (*Gelatinase Peptide)* to enable comparison of methyl sidechains as both peptide and peptoid configurations (**Figure S5A**). Interestingly, we found that the *Gelatinase Peptide* was indeed cleaved faster than the *Pan-MMP Peptide* by MMP-13, MMP-2, and MMP-9, the same MMPs that could cleave *P1 NAla* at a high rate (**Figure S5B-C**). Taken together, these substrates highlight key positions where the side chains on *N*-substituted residues may maximize differences in cleavage behavior among the set of selected MMPs.

Finally, the similarity scan library was investigated using CD (**Figure 5**) to evaluate how peptoid substitutions influenced the higher order structure of the substrates. The *P3 NPro* and *P3 Npea* substrates exhibited the most substantial deviation in structure by CD from the *Pan-MMP Peptide*. Other substitutions, especially sarcosine, gave rise to little difference in CD spectra, indicating individual peptoid substitutions do not significantly disrupt the structure of the consensus sequence. This behavior is contrasted with the full peptoid, which has no CD signal (**Figure S6**). Thus, it is expected that only the *P3 NPro* and *P3 Npea* peptomers would be resistant to proteolysis by a conformational change.

**Figure 5:**
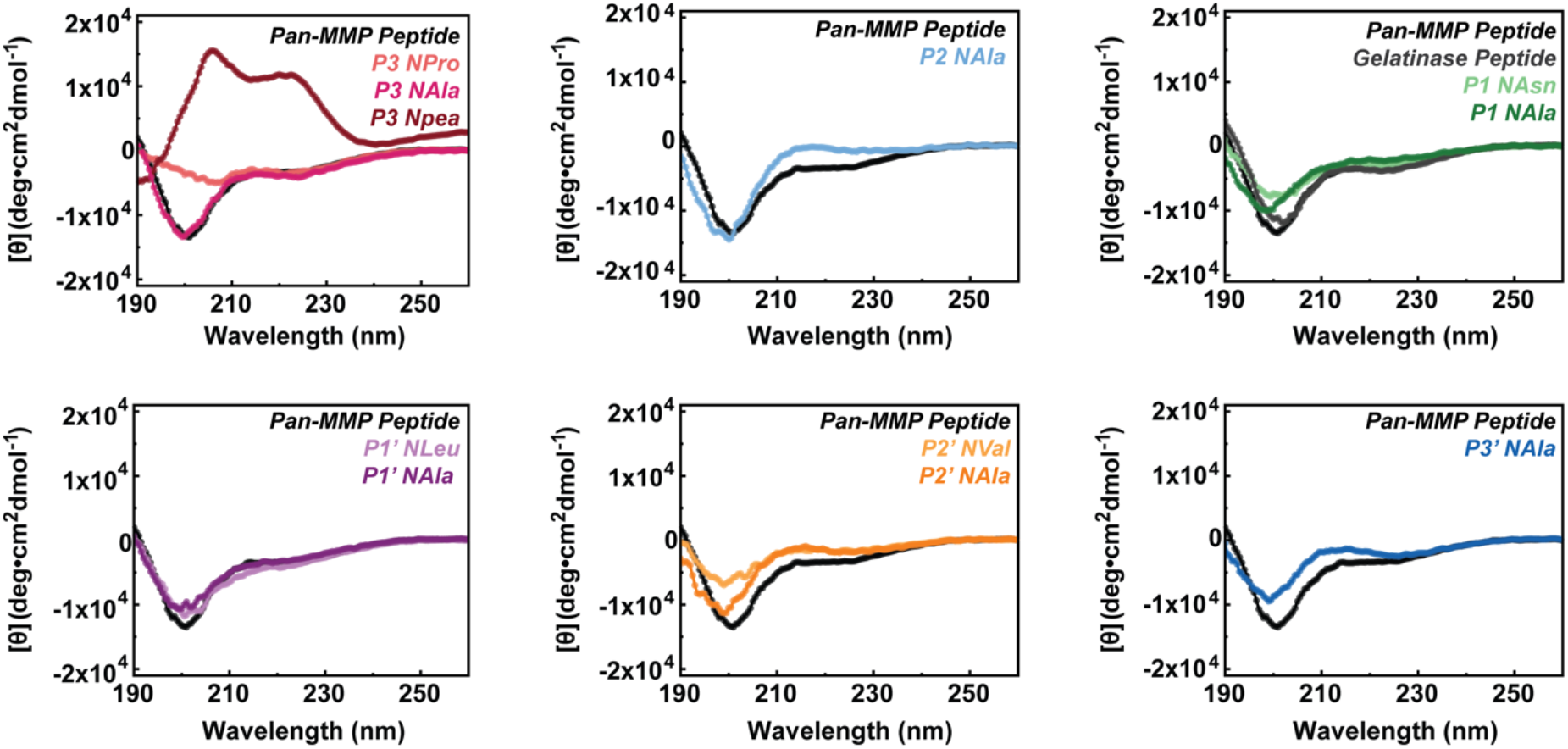
Circular dichroism spectra by substitution position. CD spectra of mean residue ellipticity shows substrates with a single peptoid substitution have little deviation in higher order structure from the *Pan-MMP Peptide* with the exception of *P3 NPro* and *P3 Npea*.

In summary, the similarity scan library matched trends seen in the peptoid analog library and demonstrated that varying sidechain structure does influence cleavage rate. We confirmed the importance of substitution site in proteolytic susceptibility, with the P1’ and P2’ sites as critical for hydrogen bonding and P1 site as non-critical for both peptide and peptoid residues. The versatility of this position may be valuable for full position scans in future combinatorial libraries.

### Combined N-substitutions for controlled degradation rate

Having identified which substrate positions are critical to proteolysis, we next hypothesized that increasing the number of *N*-substitutions would provide an additional handle for maximizing the differential response of the MMPs. Thus, we examined the impact of combining peptoid substitutions in the P3, P1, and P3’ sites. We made three tandem substituted substrates, each with two substitutions: *P1 NAsn P3’ NAla, P3 NAla P1 NAsn*, and *P3 NAla P1 NAla* (**Figure 6A, Figure S7A**). We sought to compare their rates of cleavage and degradation products to the *Pan-MMP Peptide*, as well as to the peptomers with corresponding individual substitutions. We expected that combining peptoid substitutions would further decrease the cleavage rate relative to the individually substituted peptomers by further disrupting the molecular conformation needed for enzyme recognition. Comparing amount of hydrolysis for the tandem substituted substrates to their constituent peptomers (**Figure 6B, Figure S7B-C**), we verified that they behaved in this predicted manner: after 3h, *P1 NAsn P3’ NAla* had hydrolyzed to a lower extent for all MMPs than either *P1 NAsn or P3’ NAla*. The *P3 NAla P1 NAsn* and *P3 NAla P1 NAla* substrates exhibited similar behavior, although they both hydrolyzed to a greater degree for all MMPs than *P1 NAsn P3’ NAla*. This logical outcome provides a straightforward method for engineering the degradation rate of biopolymers. Furthermore, the degradation of the tandem substituted substrates depends on substitution position, thereby enhancing the variance achieved within this small library.

**Figure 6:**
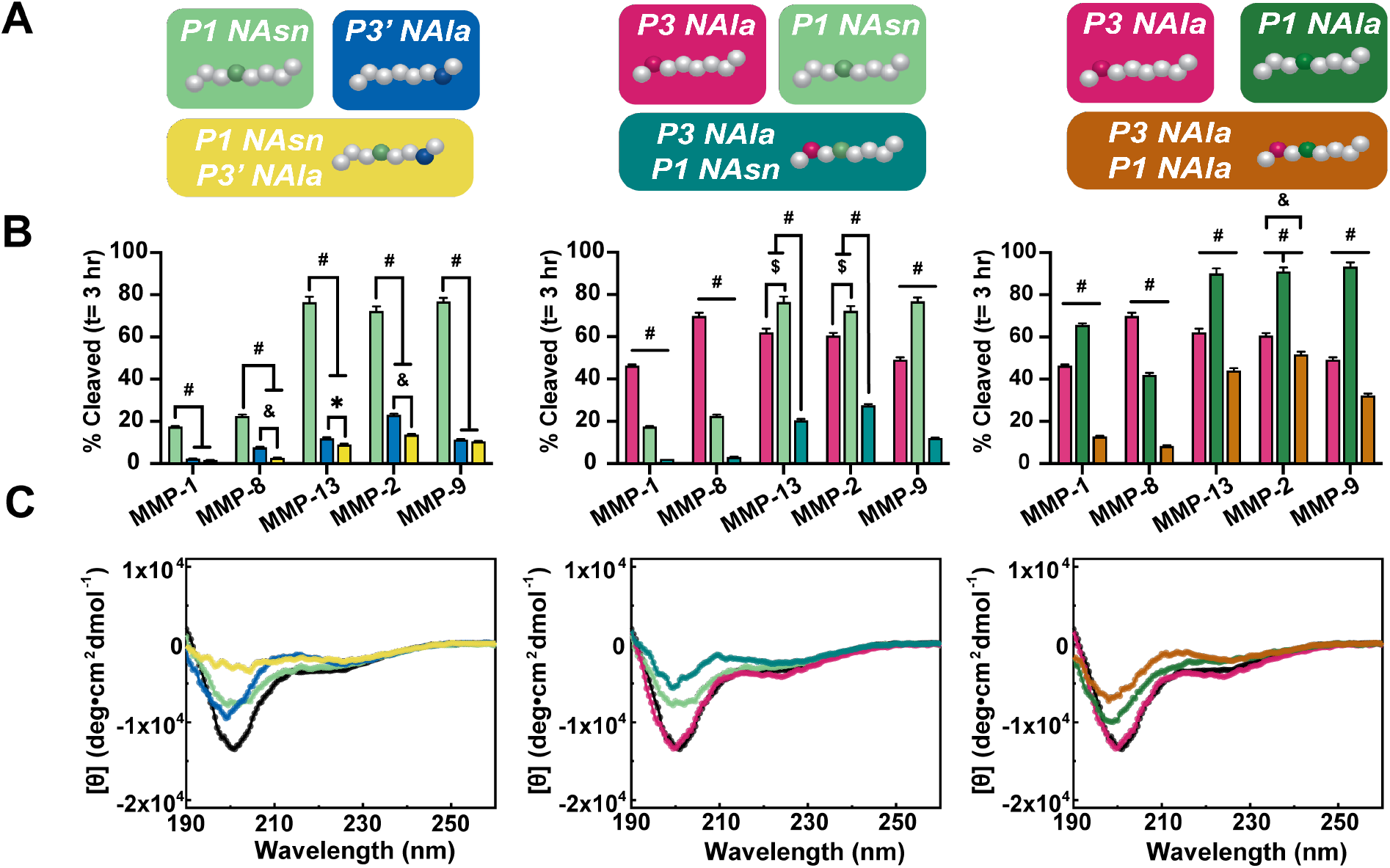
Tandem substitutions library. (A) Cartoon illustrations of individual substitutions combined to into three tandem substituted substrates. (B) Percent of substrate hydrolyzed after three hours. Error bars represent the standard error of the mean. Symbols represent statistical difference between rates: # = p<0.0001, $ = p<0.001, & = p<0.01, and *= p<0.05. (C) CD spectra of mean residue ellipticity comparing the tandem substituted peptomters to their constituent peptomers and the *Pan-MMP Peptide*.

To better understand the mechanism of this tunability we again turned to CD in search of correlations between peptomer structure and susceptibility to proteolysis. We examined the structure of the substrates with CD and saw a dampening of higher order structure roughly proportional to the combination of the individual substitutions, wherein the P1/P3’ combination caused a greater disruption than the P3/P1 combination (**Figure 6C**). The *P1 NAsn P3’ NAla* substrate had little CD signal, whereas the P3/P1 substrates retained the shape of the curve for the other substrates with decreased amplitude. Since *N*-substitution produces achiral residues, it is reasonable to presume that having substitutions on both the prime and non-prime motif would disrupt higher order structure more significantly than those contained to one side of the substrate. The tandem substrates’ deviation in structure from that of the *Pan-MMP Peptide* does appear to correlate with their proteolytic susceptibility (**Figure S8**). Given that the *P3 NAla P1 NAsn* and *P3 NAla P1 NAla* substrates have peptoids in the same positions, however, it is clear structure is not the only determinant of substrate recognition. Previously published work on the proteolytic susceptibility of *N*-methylated amino acid substitutions postulated that *N*-methylation disrupted individual enzyme-substrate contacts, rather than shifting the ensemble conformer into a form unrecognizable by proteases.^65^

We believe our peptoid-substituted substrates are influencing enzyme affinity in a similar manner. Thus, tandem substitutions result in compounding effects to both structure and proteolytic susceptibility.

### Kinetic parameters inform MMP-peptomer interactions

The three libraries of peptomer substrates investigated revealed trends in susceptibility to MMPs with peptoid location and sidechain identity. Through circular dichroism and molecular mapping of binding interactions expected in MMP-substrate complexes, we have begun to understand the mechanistic underpinnings of these trends. Specifically, we believe that the lack of hydrogen bonding donors drives the inhibition of cleavage when sidechains are translocated in the P1’ and P2’ positions. In the P1 substrate position, catalytic conversion is dictated by the steric interactions of the sidechain for both peptide and peptoid residues. Beyond these substrate positions, the affinity for the substrate may be influenced with peptoid substitutions by conformational changes or other binding factors, especially when multiple peptoid residues are incorporated. To evaluate the validity of these hypotheses, we sought to determine kinetic parameters that describe the relation between peptomer substrates and MMPs (*i*.*e*., Michaelis constant (K_M_), turnover number (k_cat_), and catalytic efficiency).

Due to material limitations, determining these parameters using traditional initial velocity analysis was infeasible, and we instead opted for a combination of progress curve assays that enabled parameter approximation directly from a reaction curve of product formed over time. First, we employed a Bayesian inference approach^54^ based on the total quasi-steady-state model (tQSSM) of differential rate equations describing the catalytic reaction.^66^ Importantly, this sophisticated approach has little bias for enzyme and substrate concentrations and does not require prior knowledge of K_M_ as many traditional methods do. First, MMPs were active site titrated at their activity-normalized concentrations to determine accurate molar concentrations for kinetic parameter fitting (**Figure S9**). Next, in addition to the screenings with 10 μM of substrate already conducted, a subset of key substrates were also screened at 0.5 μM. The fluorescence curves were then converted to progress curves by normalizing by the fluorescence saturation corresponding to complete cleavage (**Figure S10**). Combining datasets at two different concentrations enabled more accurate estimation of kinetic constants with the tQSSM (**Table S2**). However, this method is best suited for full progress curves where the substrate is 100% cleaved, a condition not satisfied for multiple substrates after the three hours of degradation tracking. Thus, the 0.5 μM screenings also provided data for the second approach used to ascertain the catalytic efficiency, the low substrate concentration approximation (**Table S3**). When substrate concentration is well below K_M_ (a condition verified to be true at 0.5 μM) the catalytic reaction is effectively first-order in rate, providing an estimation of the proportion of k_cat_ to K_M_.^55,56^

The catalytic efficiencies calculated from the two approaches showed good agreement (**Table 1, Table S4**). Catalytic efficiencies calculated using the tQSSM for the *Pan-MMP Peptide* spanned from 34,000 M^-1^ s^-1^ for MMP-1 to 1,700,000 M^-1^ s^-1^ for MMP-2 (**Table S2**), emphasizing the different activity levels of the MMPs and matching trends of relative MMP activity from other studies.^44,67,68^ When comparing the peptomers to the peptides, peptoid substitutions had a greater impact on the catalytic efficiencies for MMP-1 and MMP-8 than the other MMPs, with the cleavage of *P3 NAla* by MMP-8 as a notable exception (**Table S4**). This result is especially apparent for the P1 substituted substrates, which maintained over 60% of catalytic efficiency as compared to the *Pan-MMP Peptide* for substrates cleaved by MMP-13, MMP-2, and MMP-9, but diminished to 5% or less with MMP-1 and MMP-8. Indeed, the catalytic efficiencies of the two P1 substituted peptomers are within an order of magnitude of the peptides for MMP-13, MMP-2, and MMP-9. In general, the P1 peptomers have K_M_ values that are very similar to those of the peptides, and the changes in catalytic efficiency are driven by changes to the turnover number (**Table S2**). This is contrasted by P3, P3’, and tandem substituted peptomers, which exhibit significant increases in K_M_. This increase represents lower affinity of the enzyme for the substrate, aligning with our hypothesis that substitutions in these positions can cause conformational changes that impact enzyme recognition and binding.

**Table 1:** Kinetic parameter estimates for MMP-2. Kinetic parameters calculated using the tQSSM and the low substrate concentration approximation for catalytic efficiency. Kinetic parameters for the other MMPs are included in **Tables S2-S4**.

| Substrate | tQSSM Approximation |  |  |  |  |  | Low Substrate Concentration Approximation |  |  |  |
| --- | --- | --- | --- | --- | --- | --- | --- | --- | --- | --- |
| | $K_M$ ($\mu\text{M}$) | $K_M$ SD | $k_{\text{cat}}$ ($\text{s}^{-1}$) | $k_{\text{cat}}$ SD | Catalytic Efficiency ($\text{M}^{-1} \text{s}^{-1}$) | Catalytic Efficiency SD | $k_{\text{obs}}$ ($\text{s}^{-1}$) | $k_{\text{obs}}$ SE | Catalytic Efficiency ($\text{M}^{-1} \text{s}^{-1}$) | Catalytic Efficiency SE |
| Pan-MMP Peptide | 10 | $\pm 2$ | 17 | $\pm 2$ | 1,700,000 | $\pm 300,000$ | 6.63E-04 | 3E-06 | 1,540,000 | $\pm 70,000$ |
| Gelatinase Peptide | 21 | $\pm 6$ | 60 | $\pm 10$ | 2,600,000 | $\pm 900,000$ | 1.30E-03 | 1E-05 | 3,000,000 | $\pm 100,000$ |
| P3 NAla | 130 | $\pm 30$ | 50 | $\pm 10$ | 400,000 | $\pm 100,000$ | 1.60E-04 | 1E-06 | 370,000 | $\pm 20,000$ |
| P1 NAsn | 18 | $\pm 3$ | 14 | $\pm 2$ | 800,000 | $\pm 200,000$ | 4.03E-04 | 2E-06 | 940,000 | $\pm 40,000$ |
| P1 NAla | 12 | $\pm 2$ | 22 | $\pm 3$ | 1,800,000 | $\pm 400,000$ | 7.62E-04 | 5E-06 | 1,770,000 | $\pm 80,000$ |
| P3' NAla | 50 | $\pm 20$ | 4 | $\pm 1$ | 90,000 | $\pm 50,000$ | 5.04E-05 | 5E-07 | 117,000 | $\pm 6,000$ |
| P1 NAsn P3' NAla | 200 | $\pm 100$ | 8 | $\pm 5$ | 50,000 | $\pm 40,000$ | 3.28E-05 | 6E-07 | 76,000 | $\pm 4,000$ |
| P3 NAla P1 NAsn | 110 | $\pm 30$ | 12 | $\pm 3$ | 110,000 | $\pm 40,000$ | 4.95E-05 | 4E-07 | 115,000 | $\pm 5,000$ |
| P3 NAla P1 NAla | 200 | $\pm 100$ | 60 | $\pm 30$ | 300,000 | $\pm 200,000$ | 1.261E-04 | 7E-07 | 290,000 | $\pm 10,000$ |

### Array-based sensing for MMP differentiation

Determining kinetic parameters confirmed that, in general, the five MMPs studied responded similarly to each peptomer, making it difficult to find distinguishing features that would garner selectivity to individual MMPs. Despite recurring trends in relative activity, however, each peptomer-MMP combination produces a unique rate of hydrolysis, which may be useful for applications in array-based sensing.^69^ To visualize this variance, the three-hour fluorescent traces were fit directly to an exponential plateau function (**Figure S11**). Excluding substrates that did not cleave, we then compiled the logarithm of the k constants used to fit the exponentials (**Table S5**) and visualized each substrate-MMP relationship in a heatmap (**Figure 7A**). Importantly, this collection of differential responses reveals a specific pattern for each MMP, with log(k) values being the distinguishing features. Similar arrays of cross-reactive binding interactions have previously been used as fluorescent “fingerprints” able to differentiate between protease types.^70,71^ Here, we sought to explore the discriminatory power of a dynamic stimulus (*i*.*e*., degradation) with our tunable peptomer array using multivariate data analysis.

**Figure 7:**
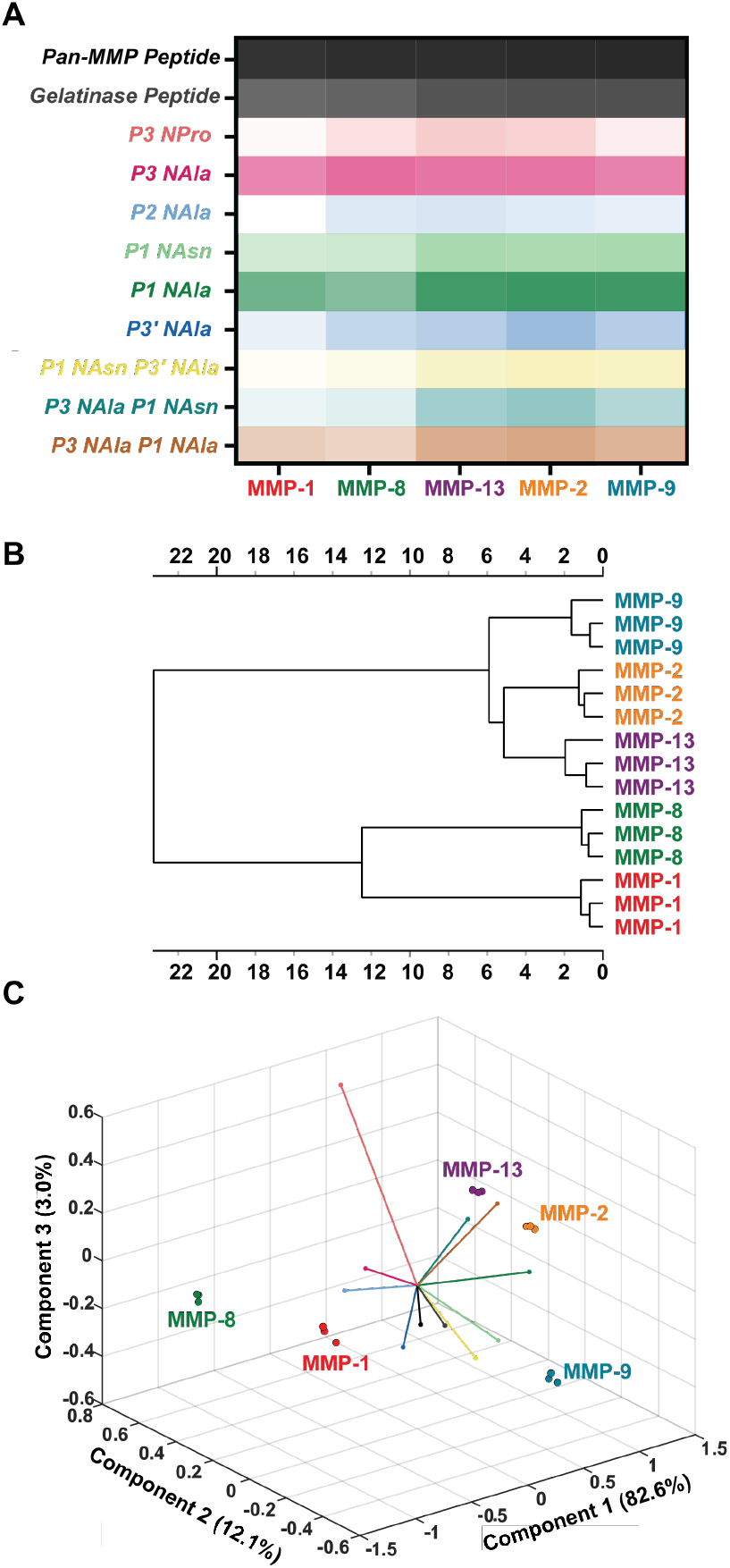
Multivariate data analysis of peptomer degradation patterns. (A) Heat map generated from log(k) values of exponential plateau fits of three-hour fluorescence traces. The peptomers are features with values that form an array, giving each MMP a “fingerprint” of degradative behavior. The heat map is colored according to the average k-constant value of the three replicates for each enzyme-substrate interaction. (B) HCA dendrogram depicting clustering of MMPs. The clustering depicts that MMP-13’s degradation patterns more closely resemble the gelatinases than the collagenases. (C) PCA biplot showing clustering of MMPs by principal component scores and loadings from each peptomer.

First, the data were subjected to unsupervised techniques to identify data that naturally cluster together (HCA) and identify the elements of the array that best separate the data (PCA),^72^ while withholding class information (*i*.*e*., MMP identity). HCA (**Figure 7B**) revealed interesting clustering of the MMPs. Given the MMPs tested belong to two unique subclasses, we expected the collagenases and gelatinases to be clustered together. Instead, the collagenases, MMP-1 and MMP-8, are separated from MMP-2, MMP-9, and MMP-13, the final of which is also a collagenase. Furthermore, the clustering revealed more similarity between MMP-2 and MMP-13 than MMP-2 and MMP-9, the two gelatinases. This outcome matches correlations observed in native peptide sequences and is an important consideration for researchers targeting the gelatinase MMPs.^19^

Next, we used PCA to evaluate the utility of the array for separation by MMP type, and to better understand the contribution of each peptomer towards clustering. PCA seeks to project the data in a low dimensional space using eigenvalue decomposition, resulting in *n*-dimensional axes that explain the majority of the dataset’s variance. The resulting eigenvalues produce coordinate positions for each individual sample, such that the correlation of variance of one sample to another can be evaluated.^73^ Excitingly, PCA scree plots show 97.7% captured variance with just three principal components (**Figure S12**), and revealed successful clustering and separation by MMP type for the replicates of each degradation profile **(Figure 7C, Figure S13)**. Importantly, because PCA places no bias on finding the greatest variance between samples, this means that the replicates of the same MMPs are treated identically, that is, as distinctly different samples. Therefore, clustering of the samples in PCA means that the variance between MMP replicates is indeed smaller than the variance between different MMPs, and confirms that peptomers behave in a sufficiently unique manner for use in differential sensing applications.

To better understand how the response of each peptomer contributes to the separation of the data, a loading plot was generated and superimposed onto the PCA score plot **(Figure 7C)**. Loading plots illustrate the influence each original variable contributes to each principal component axis, in which the vector magnitude is directly proportional to the eigenvector weights, with a higher weight correlating to more influence of the original variable.^74^ Thus, the proximity of each peptomer vector’s endpoint to a data point reveals whether that variable is important for discriminating that particular MMP. The *Pan-MMP Peptide* vector is short because the enzyme concentrations were normalized to cleave this substrate at the same rate. Thus, it was included for reference rather than its discriminatory ability. Briefly, the *P3 NPro* and *P2 NAla* substrates are important in clustering MMP-1 and MMP-8, demonstrating that substrates with low hydrolytic susceptibility still provide powerful information. On the other side, the P1 and P3/P1 tandem substituted substrates help separate MMP-2, MMP-9, and MMP-13, despite their overlap in specificity. Overall, the biplot (score plot + loading plot) reveals that the array of peptomers is valuable and informative in acquiring characteristic and diverse responses to discriminate the proteases.

Finally, we investigated the potential for this type of data to generate a model that could classify MMPs from an unknown sample. Following unsupervised analysis, the principal components were used to reconstruct the original data (**Table S6**). These new linear combinations were fed to a supervised pattern-recognition algorithm to train a model for future classifications. Namely, we used the kNN algorithm with three-fold cross-validation.^75^ Thus, for each fold, the algorithm is being trained on 2/3 of the replicates for each MMP and seeking to identify the remaining 1/3 without knowing the class labels. It does so by computing the distances between a testing data point and all the data points in the training set. It then classifies the unknown point by matching it to its nearest neighbors with the shortest distance. When the transformed data from two principal components were fed to the algorithm, the model was 100% accurate. Furthermore, when the raw data (log(k) constants) were fed directly to kNN, it also resulted in 100% classification accuracy (**Figure S14**). Each k-constant concisely quantifies an MMP-enzyme interaction, thus demonstrating the proficiency of this quantification. While this is a limited dataset and the degradation profiles depend on the concentration of substrate and protease, the successful clustering and classification of different MMP types demonstrates the utility of peptomers as differential array sensors for distinguishing between proteases with similar specificities.^76^ To achieve utility in diagnostic applications, the training dataset can be expanded to include mixtures of proteases and inhibitors, as modeled by Lauffenburger’s Proteolytic Activity Matrix Analysis (PrAMA), which similarly relied on the deconvolution of dynamic signals from multiple peptide-based substrates.^77^

### Controlled release of fluorescent peptomers from a hydrogel

In addition to serving as probes for array-based sensing, proteolytically degradable peptomers will be useful for biomaterial applications requiring a reliable, tunable rate of release. Peptoids offer means to modulate the proteolytic susceptibility of the substrate without making major changes to the overall chemical nature of the biopolymer, in contrast to peptide-based substrates, where swapping out an amino acid may change the overall polarity of the molecule, alter the bioactivity of the substrate, and perhaps lead to off-target proteolysis. Degradability is an important parameter to tune for hydrogels used in controlled-release and cell culture applications.^68^ To demonstrate the advantages of employing peptomers in hydrogel applications, we implemented peptomer substrates as degradable probes in a PEG hydrogel (**Figure 8A**).

**Figure 8:**
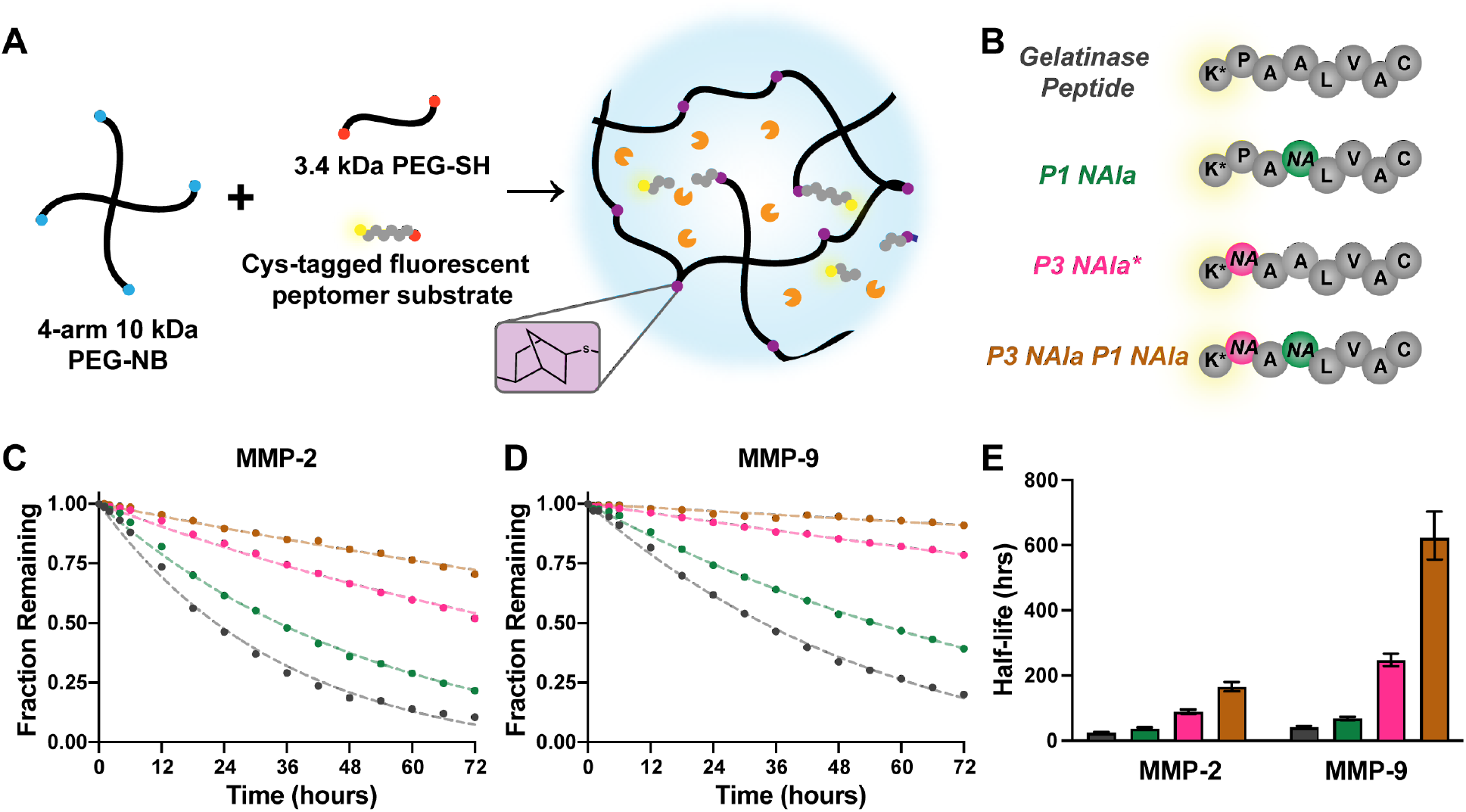
Controlled-release of peptomer substrates from hydrogel. (A) Cartoon schematic of the PEG hydrogel network crosslinking with thiol-ene chemistry. Fluorescent substrates were tethered in as dangling ends. When cleaved by an enzyme, the fragment containing the fluorophore diffuses out of the gel into the surrounding solution. (B) Controlled release substrates employed. Substrates included the *Gelatinase Peptide* sequence and three peptomers. *P3 NAla\** is slightly distinct from the *P3 NAla* substrate in previous investigations, as the substrate used here has an alanine in the P1 position versus an asparagine as the substrates based on the *Pan-MMP Peptide* had. Cysteines were included at the amino termini for thiol-ene crosslinking into the network. 7-Methoxycoumarin-4-acetic acid-modified lysine was again used as the fluorophore on the carboxy termini for fluorescent tracking. (C) Fraction of substrate remaining in the gel over 3 days for MMP-2 and (D) MMP-9. Standard deviations were calculated, but the error bars were too small to appear on the plot. (E) Half-lives of substrate release from exponential decay fits. Error bars represent the 95% confidence interval.

To tune the rate of release, we synthesized four substrates with varying degrees of degradability by MMP-2 and MMP-9. The substrates were based on the highest catalytic efficiency *Gelatinase Peptide* sequence, and they were designed with a modified-lysine fluorophore on the amine terminus and a cysteine on the carboxy terminus for thiol-ene crosslinking (**Figure 8B**). *NAla* residues were substituted in the P3, P1, and together in the P3 and P1 positions to tune cleavage rate. Gels were immersed in buffered solution containing MMP at the previously determined activity-normalized concentrations (0.43 nM for MMP-2 and 2.8 nM for MMP-9, as quantified by active site titrations). These concentrations are well below reported exogenous protease concentrations of MMP-2 and MMP-9 (20 nM and 40 nM, respectively)^12^ and bacterial collagenase (0.2 mg mL^-1^)^78^ used in other representative studies of degradable hydrogels, demonstrating the sensitivity afforded with peptomer substrates. For cell culture applications, the concentration of secreted MMPs has been estimated to be near 100 nM for human mesenchymal stem cells seeded at 0.5 × 10^5^ cells mL^-1^.^79^ This concentration includes many types of MMPs and perhaps other proteases with similar specificity; thus, we believe investigating MMP-2 and MMP-9 at lower concentrations provides useful insight to the behavior of these individual enzymes when diffusion-limited in a hydrogel.

We found that the substrates tethered to the gel degraded in the same order as the substrates in solution assays (*Gelatinase Peptide* > *P1 NAla* > *P3 NAla* > *P3 NAla P1 NAla*) for both MMPs (**Figure 8C-D**). Degradation half-lives for MMP-2 spanned a range from 24 to 166 hours. For MMP-9, half-lives were between 42 and 624 hours (**Figure 8E**). Despite using activity-normalized concentrations, MMP-2 cleaved every substrate faster than MMP-9. This is likely in part due to its smaller molecular weight, allowing it to diffuse through the network faster and cleave the substrates. MMP-2 and MMP-9 are often both dysregulated in pathological conditions^80–84^ and this experiment demonstrates how their inherent difference in activity and size garners some selectivity. Using peptomers, we are able to target certain enzymes and modulate the rate of degradation.

## Conclusion

In conclusion, we have explored the use of peptoid substitutions as residues to program distinct degradation behavior into MMP substrates. We identified straightforward design rules for incorporating peptoid substitutions into MMP-degradable substrates: 1) peptoid substitutions are particularly tolerated at the P3, P1, and P3’ substrate positions of the active site, 2) peptoid substitutions are not tolerated at the P1’ and P2’ sites due to critical hydrogen bonding interactions, and 3) combining substitution sites provides a straightforward strategy to control the rate of degradation for various applications. Specifically, we identified substrate sites for tuning degradation rates in solution over a span of minutes (peptides and P1 substrates), hours (P3 and P3/P1 substrates), and days (P3’ and P1/P3’ substrates). The tunability of degradation rates with peptomers was further employed for controlled release from a biocompatible hydrogel. We believe the generalizability of these design rules to other MMP-degradable substrates make peptoids a valuable option for modulating proteolysis. In particular, the ability to reliably tune degradation rate without significantly changing the chemical nature of the molecule is evidence of the programmability afforded with sequence-defined non-natural oligomers.^85^ Furthermore, the variance in the rate of degradation enables separation and classification by multivariate data analysis, which can be used to distinguish between MMPs despite their closely-related specificity profiles. These results build on established understanding of MMP recognition motifs and advance the utility of degradation profiling as means to differentiate between proteases.

## ASSOCIATED CONTENT

### Supporting Information

The Supporting Information is available free of charge on the ACS Publications website and includes substrate purity traces, norbornene functionalization NMR, LC-MS degradation chromatographs, additional fluorescence traces and fits, additional CD traces, kinetic parameter determinations, inputs used for multivariate data analysis, and PCA and kNN classification results.

MATLAB code is available at github.com/mariahaustin/MMP_Peptomers_DiffDegradation.

## AUTHOR INFORMATION

### Author Contributions

M.J.A, H.C.S., C.M.W, L.D.M., and A.M.R. conceived and designed the research, M.J.A., H.C.S., C.M.W., N.R.L, and J.M.C. performed experiments, A.M.R. supervised experiments. The manuscript was written through contributions of all authors. All authors have given approval to the final version of the manuscript.

### Notes

The authors declare no competing financial interest.

## Acknowledgments

This research was supported by the National Science Foundation (DMR-2046746, A.M.R.). M.J.A., H.C.S., and L.D.M. were supported through the National Science Foundation Graduate Research Fellowship (Program Award No. 000392968). We acknowledge the use of shared facilities at the University of Texas Mass Spectrometry Facility, Proteomics Facility, and the Targeted Therapeutic Drug Discovery & Development Program (CPRIT Core Facilities Support Award (grant # RP160657)). We also thank Dr. Ron Zuckerman for his reviewing and editing assistance. Graphical abstract contains an enzyme structure created with BioRender.com using the PDB 1L6J-crystal structure of human matrix metalloproteinase MMP9 (gelatinase B) published at doi.org/10.1107/s0909444902007849.

## Graphical abstract

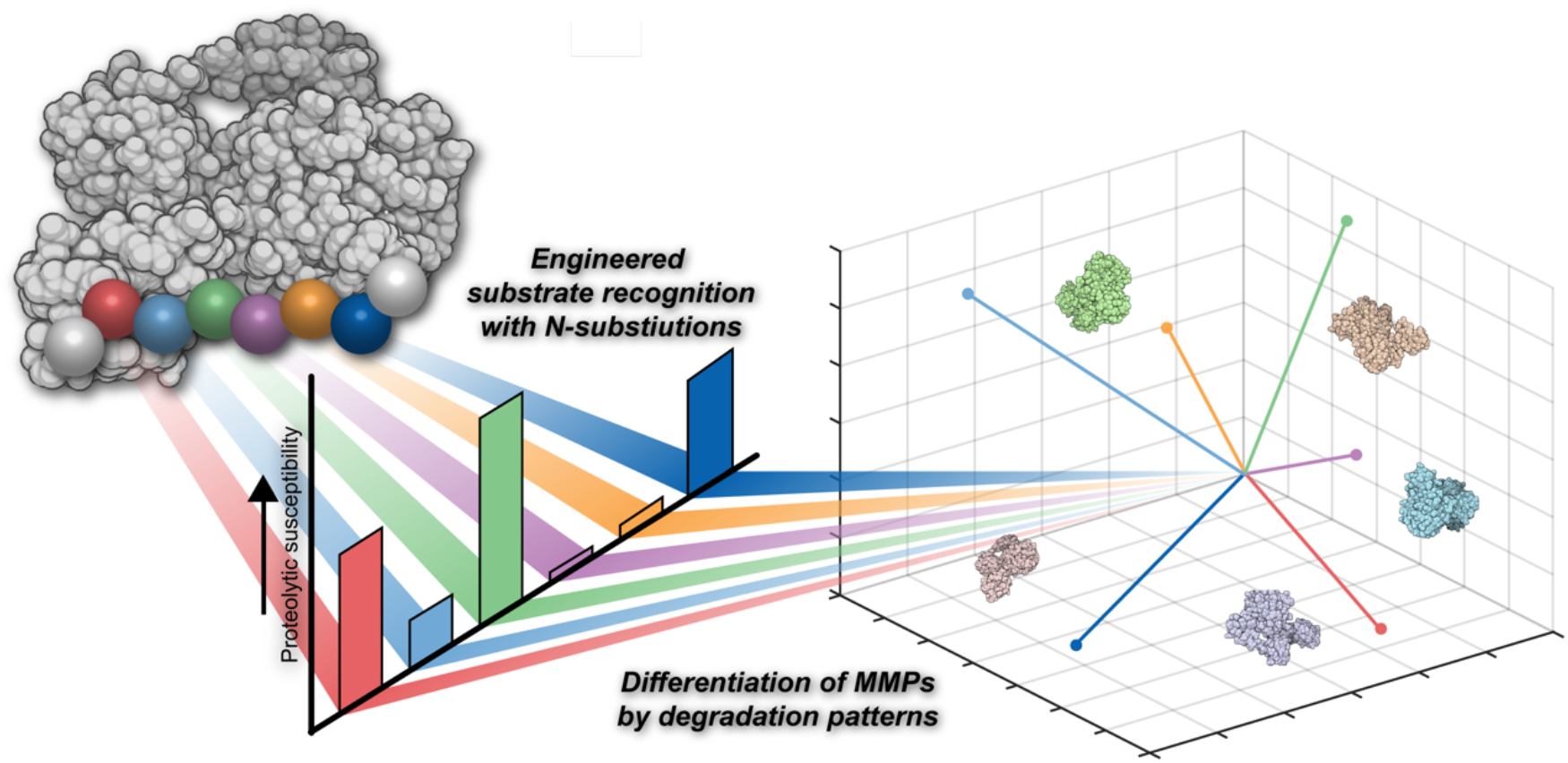

