## Supplemental Information for "Fluorescent peptomer substrates for differential degradation by metalloproteases"

**Table S1: Enzyme lot numbers.** Experiments with multiple lot numbers were screened to confirm consistency in enzyme activity.

| **Enzyme** | **Catalog Number** | **Lot Number(s)** |
|---|---|---|
| MMP-1 | 901-MP-010 | BRT0920091 |
| MMP-8 | 908-MP-010 | CLF0720101, CLF0721071 |
| MMP-13 | 511-MM-010 | CJT1220101 |
| MMP-2 | 902-MP-010 | APT0920111 |
| MMP-9 | 911-MP-010 | APU1020081, APU1121111 |
| Bacterial Collagenase | 17100017 | 2357188 |

### FRET Substrates

#### *Pan-MMP Peptide*

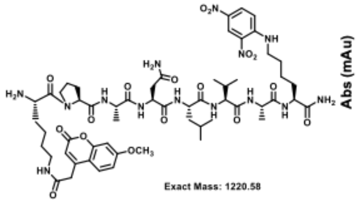

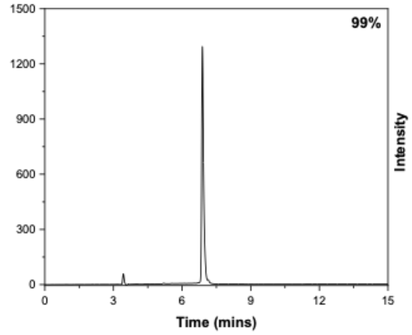

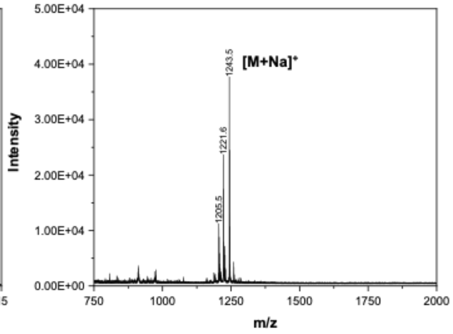

#### *Gelatinase Peptide*

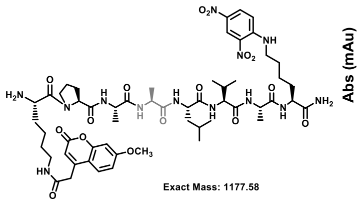

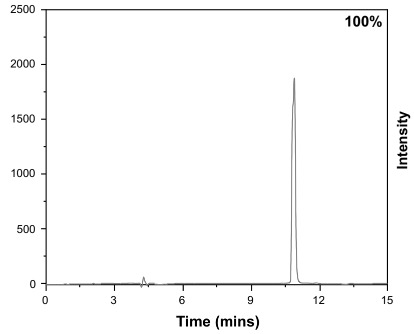

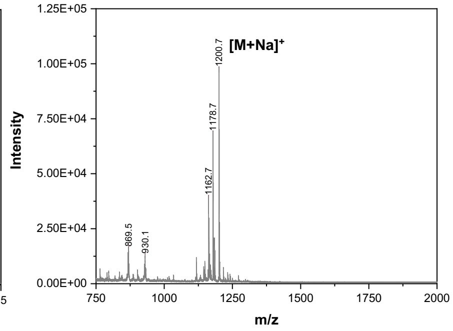

#### *P3 NPro*

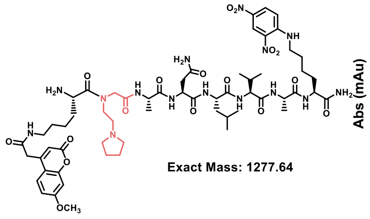

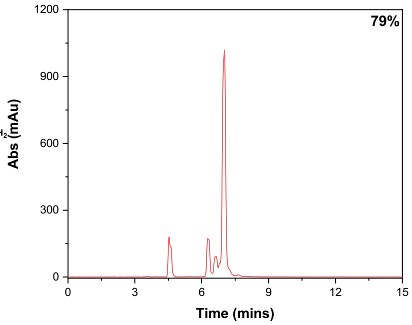

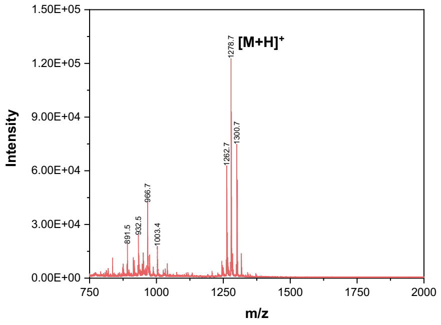

#### *P2 NAla*

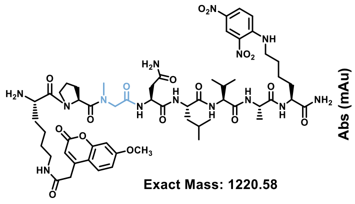

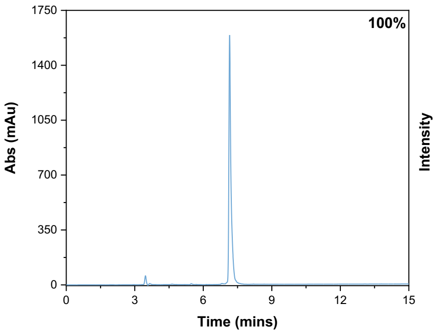

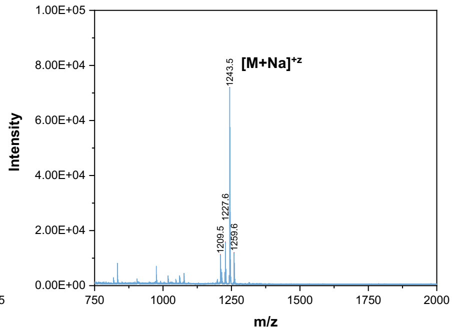

***P1 NAsn***

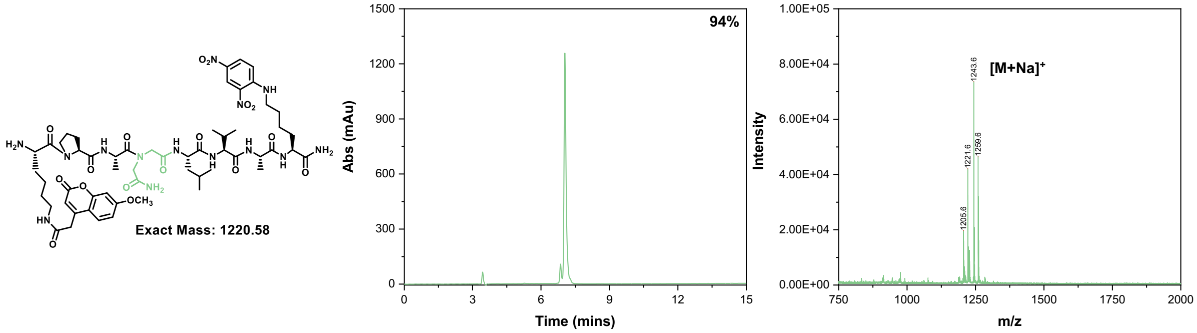

***P1' NLeu***

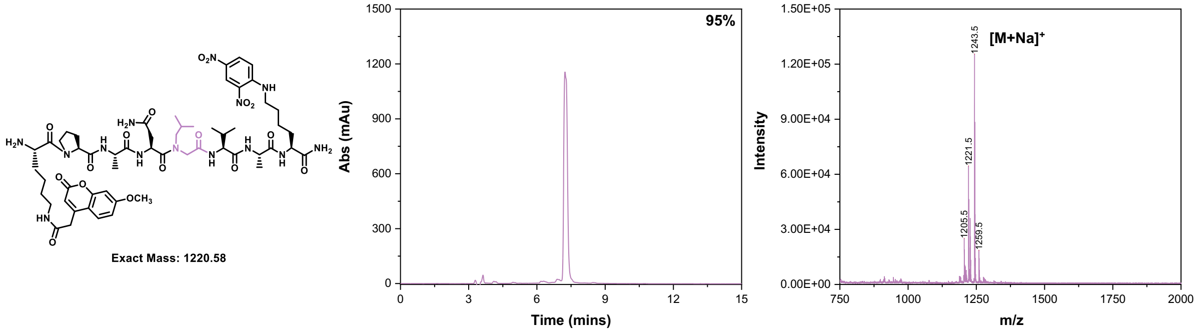

***P2' NVal***

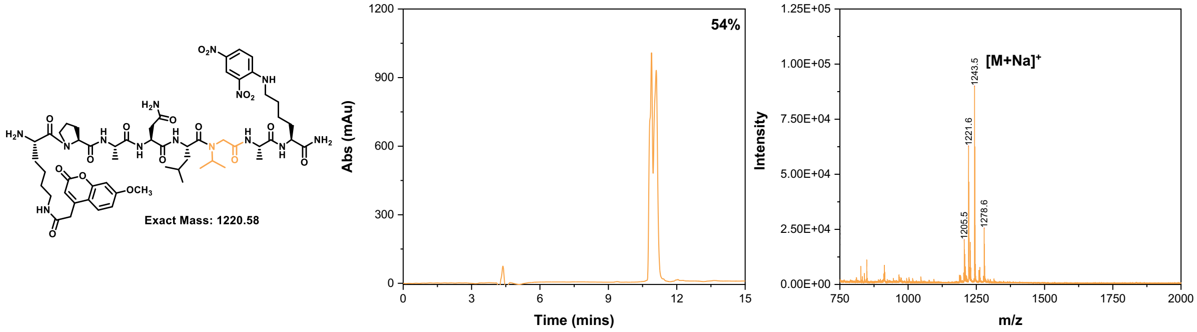

***P3' NAla***

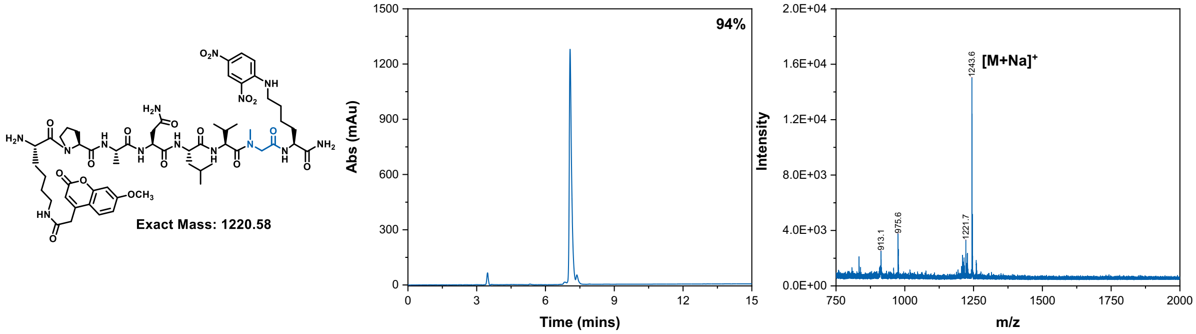

***Peptoid***

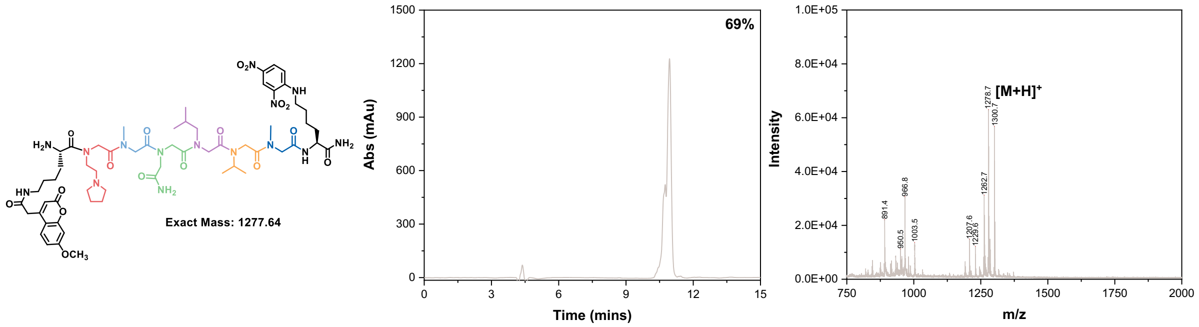

***P3 NAla***

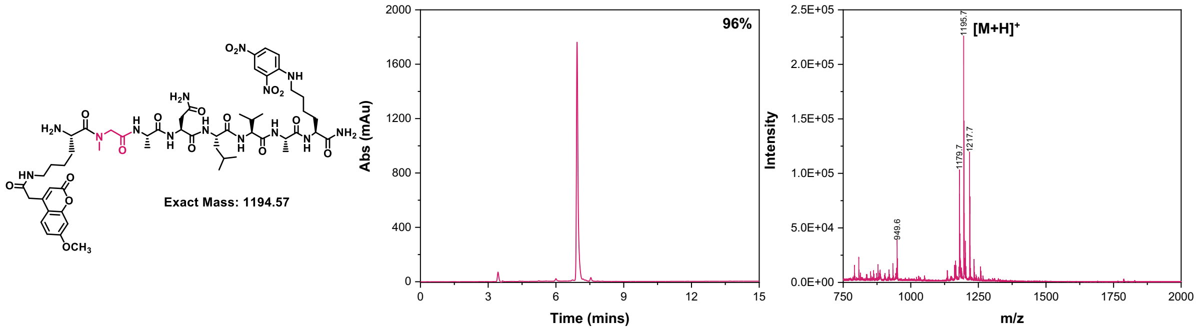

***P1 NAla***

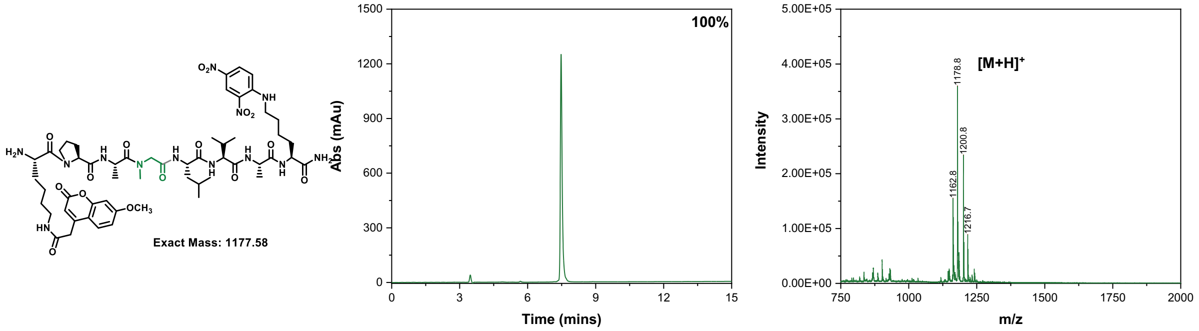

***P1’ NAla***

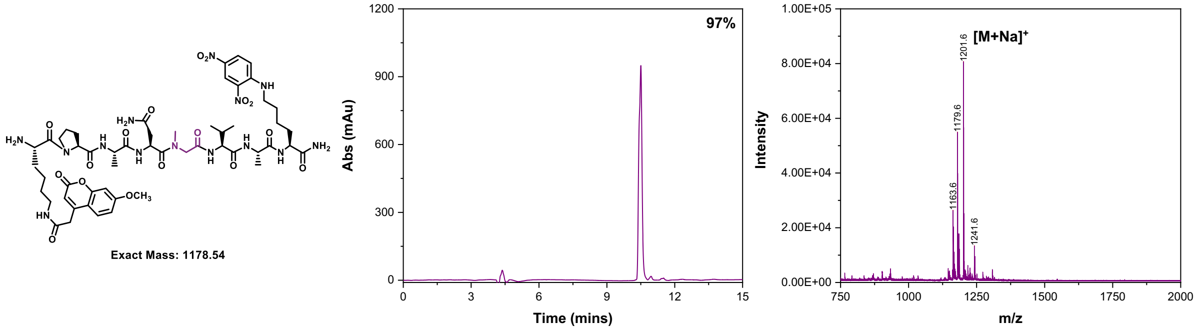

#### *P2' NAla*

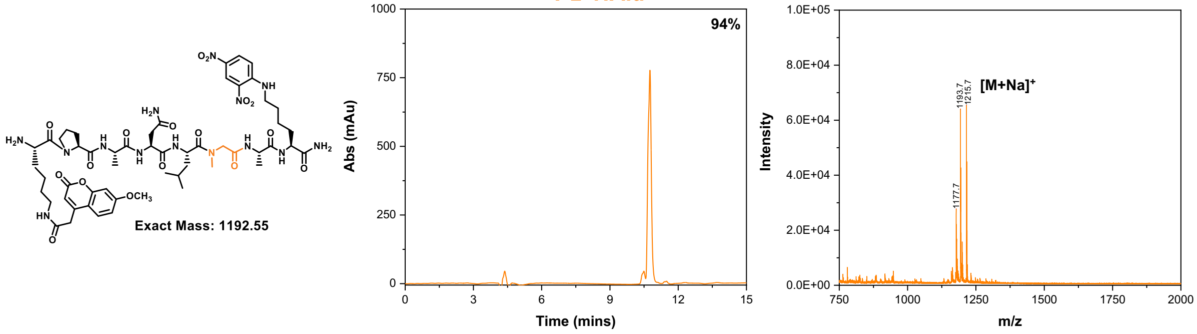

#### *P3 Npea*

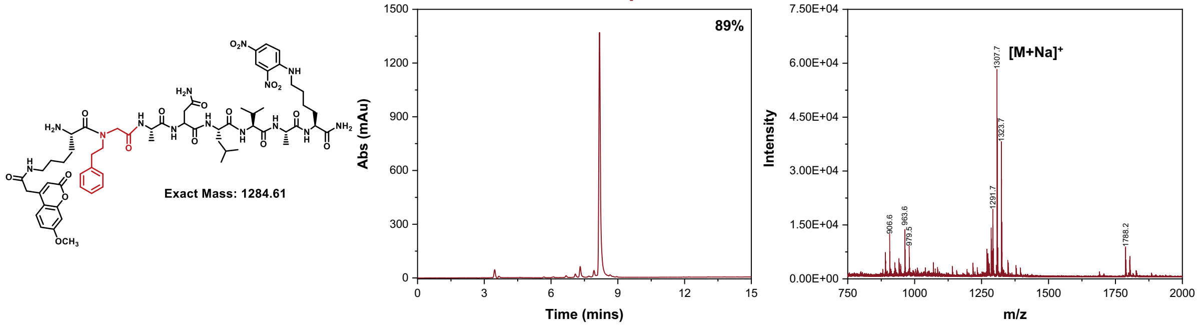

#### *P1 NAsn P3' NAla*

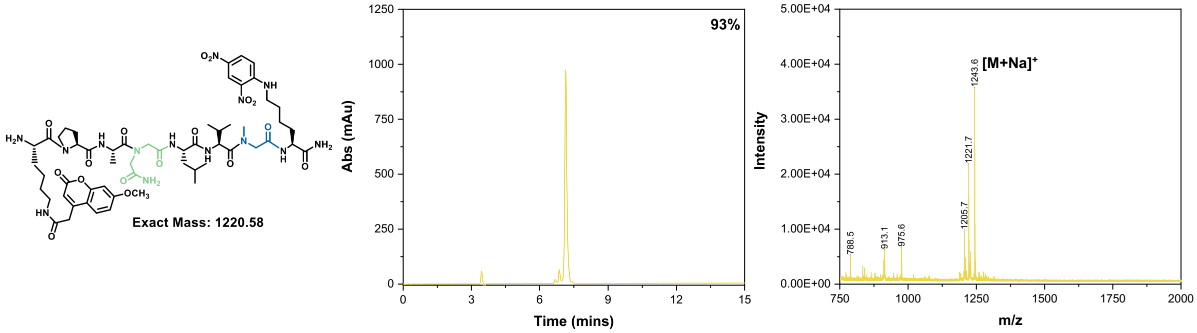

#### *P3 NAla P1 NAsn*

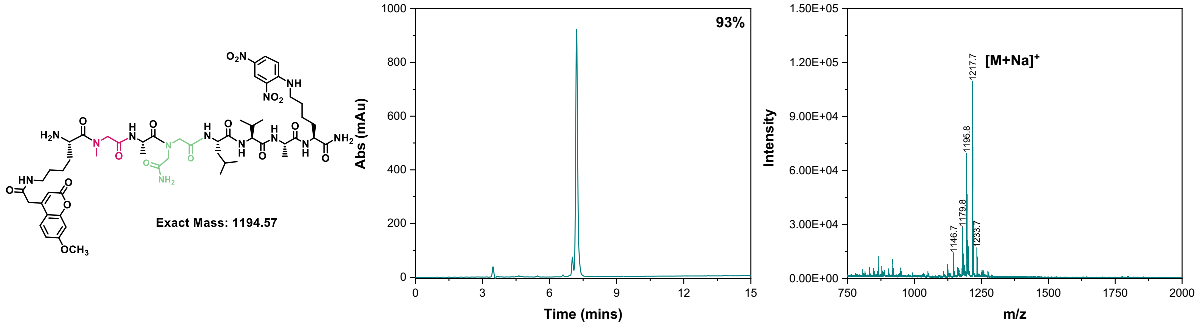

*P3 NAla P1 NAla*

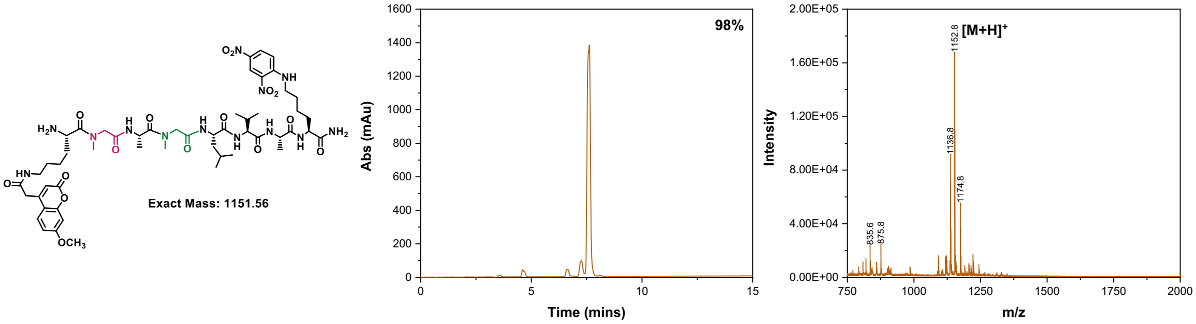

### Cysteine-modified fluorescent substrates

*Cys Gelatinase Peptide*

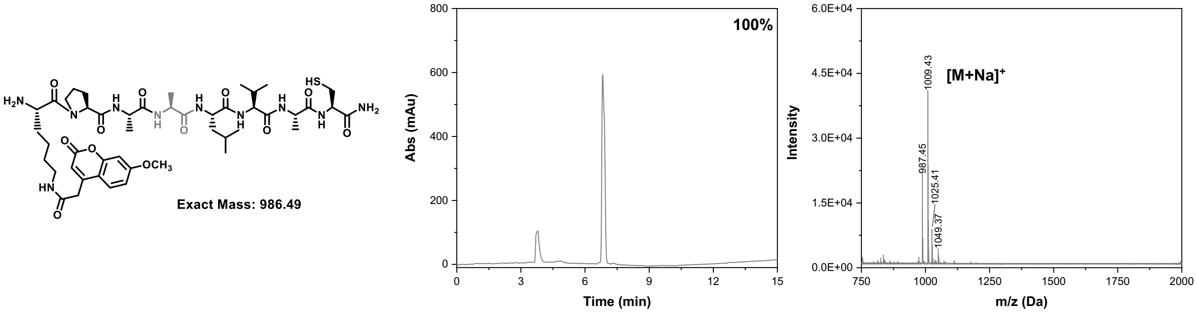

*Cys P1 NAla*

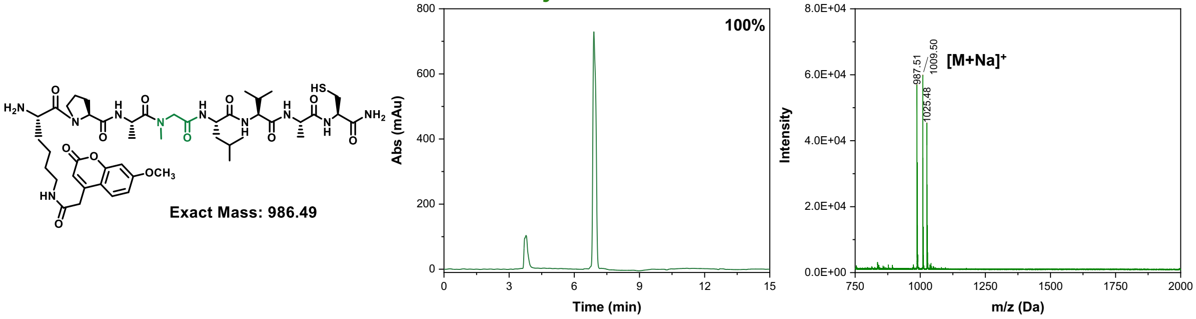

*Cys P3 NAla**

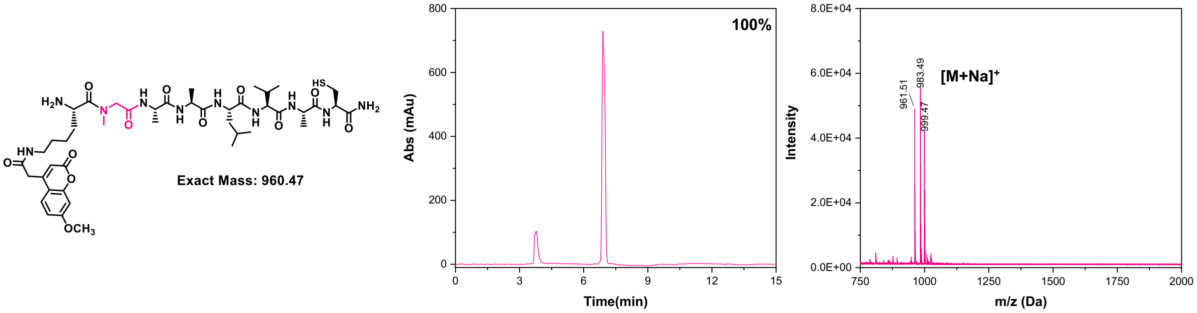

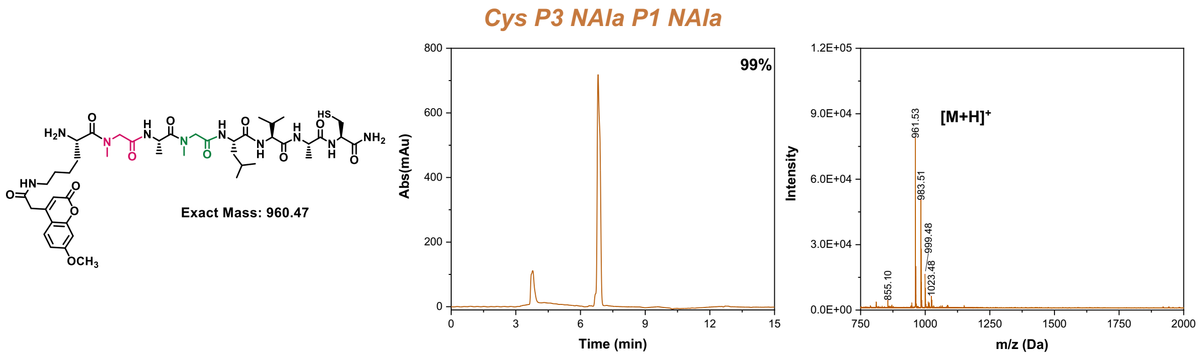

**<u>Figure S1</u>: Substrate purity traces.** Chemical structures with colored peptoid substitution and expected mass (left), analytical HPLC traces with purity determined by integration (center), and MALDI spectra to confirm molecular weight (right). HPLC trace have a solvent peak between 3-4 minutes that was excluded from purity determinations.

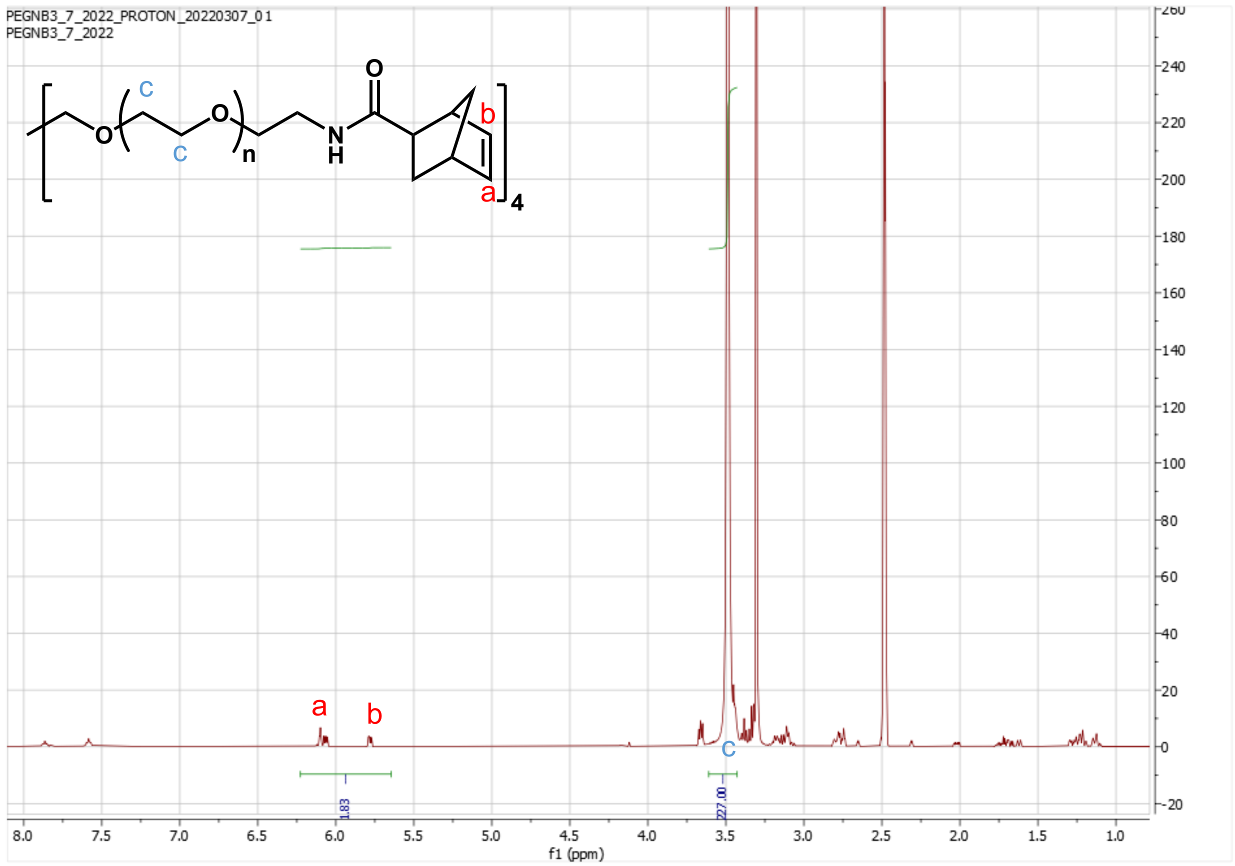

**<u>Figure S2</u>: PEG-NB functionalization determination by $^1$H NMR.** The ratio of backbone hydrogens to end group hydrogens would be 227:2 for 100% functionalization. Integration shows that the observed ratio achieved was 227:1.83, yielding 91.5% functionalization.

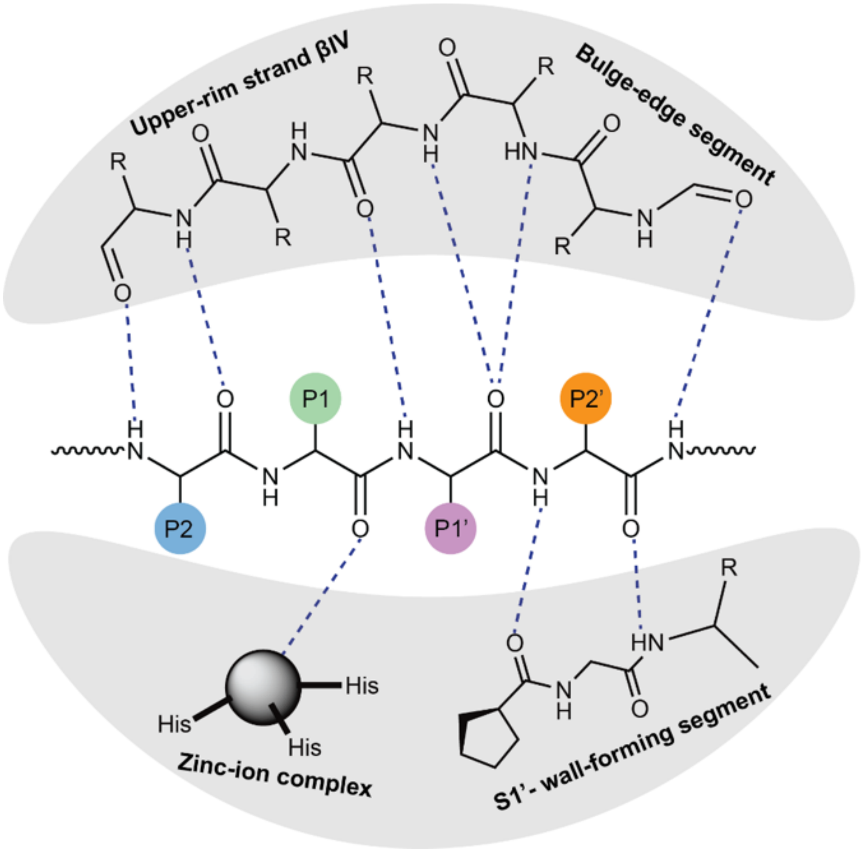

**Figure S3: Michaelis complex formed between MMP active site and substrates.** This proposed mechanism demonstrates the importance of the nitrogen-affixed hydrogens in on the prime motif and suggests that the identity of sidechains in the active site pocket influence enzyme recognition sterically. Figure adapted from ref.1 with permission from Elsevier (License number 5256041381485).

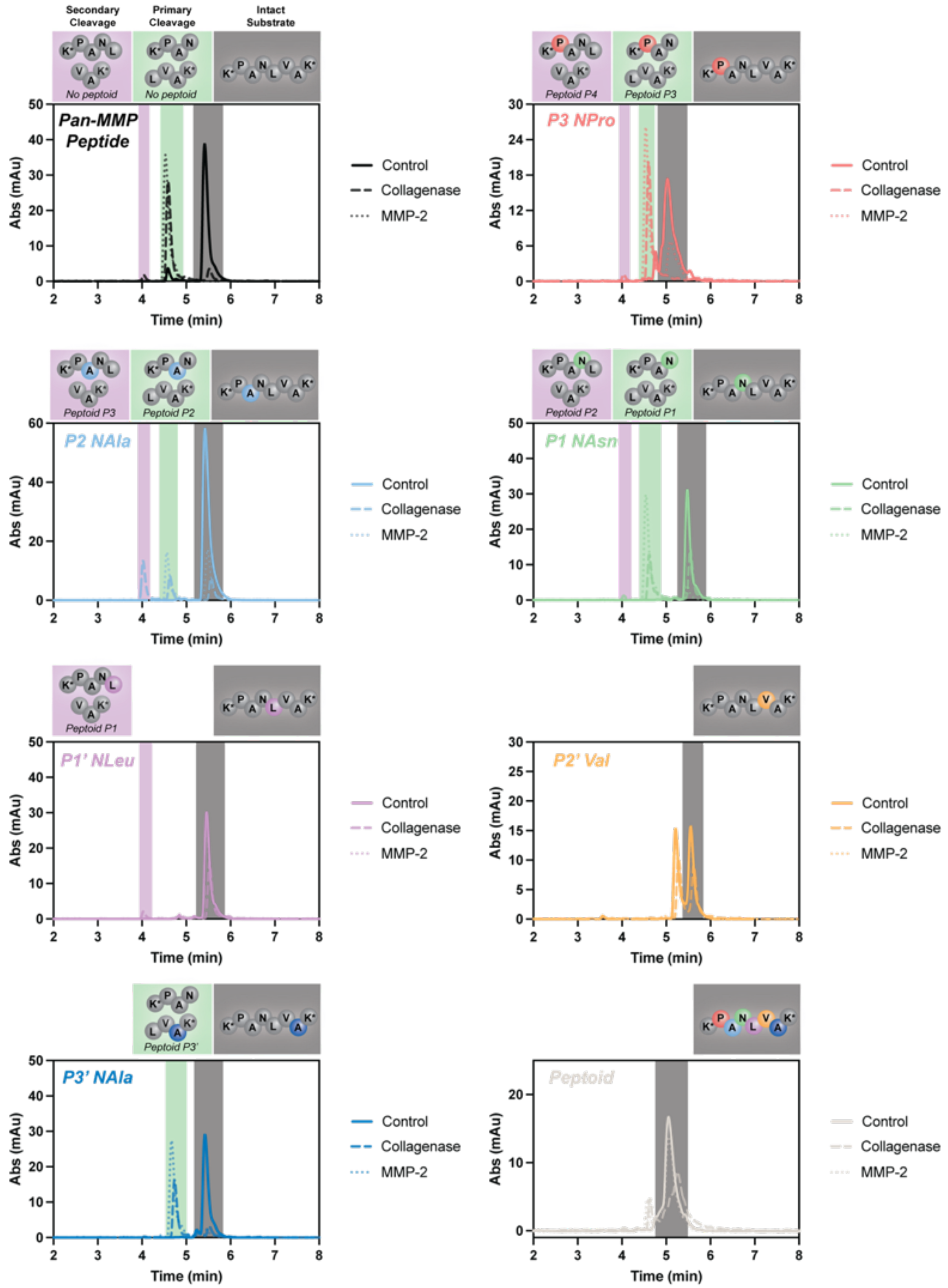

**Figure S4: LC-MS chromatographs of peptoid analog library substrates.** Traces collected after 24 hours of exposure to either buffer (control), MMP-2, or bacterial collagenase. Traces were collected at 400 nm to detect the absorbance of the dinitrophenyl quencher. Thus, there was only one peak corresponding to a single fragment from each cleavage event. Those peaks were matched to corresponding extracted ion chromatographs generated at specific m/z ratios according to the expected mass fragments to determine their identity and then integrated to determine composition of degradation components.

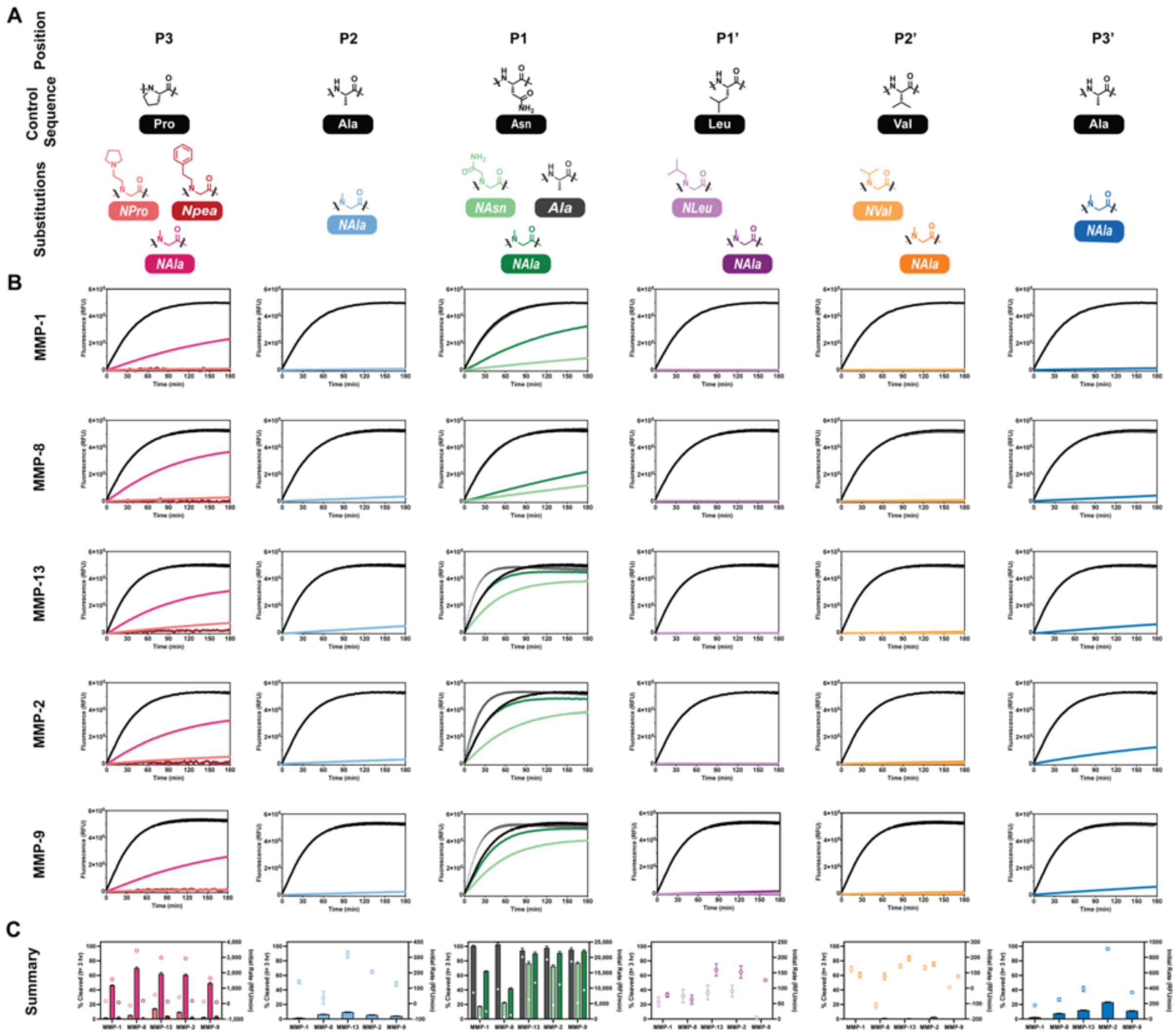

**Figure S5: Similarity scan fluorescence traces and summary plots.** (A) Consensus sequence of the *Collagenase Peptide* along with substituted residues at each position (B) Fluorescence curves measuring cleavage of each substrate over the course of three hours of exposure to MMPs. (C) Comparison of percent hydrolysis after three hours (bars, left axes) and initial rate of cleavage (unfilled circles, right axes) by residue position.

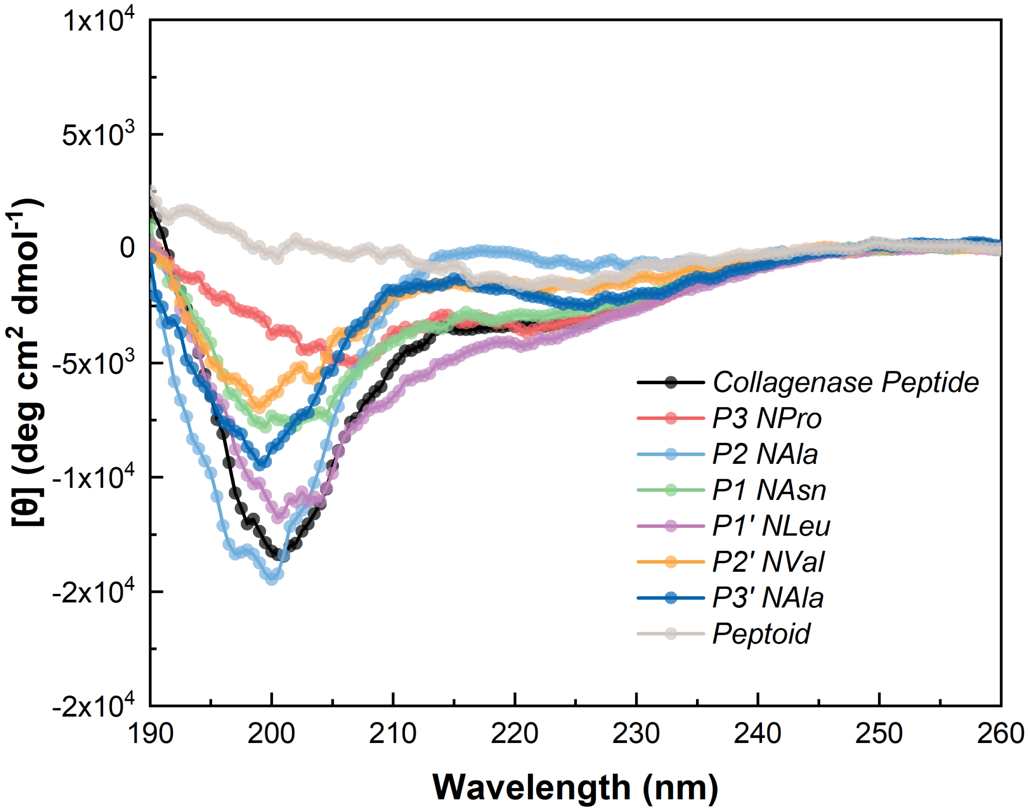

**<u>Figure S6</u>: Peptoid analog library circular dichroism.** Single peptoid substitutions cause slight disruptions in higher order structure depending on location and the *Peptoid* has no measurable structure, as expected with removal of all hydrogen bonding locations in the active sequence backbone.

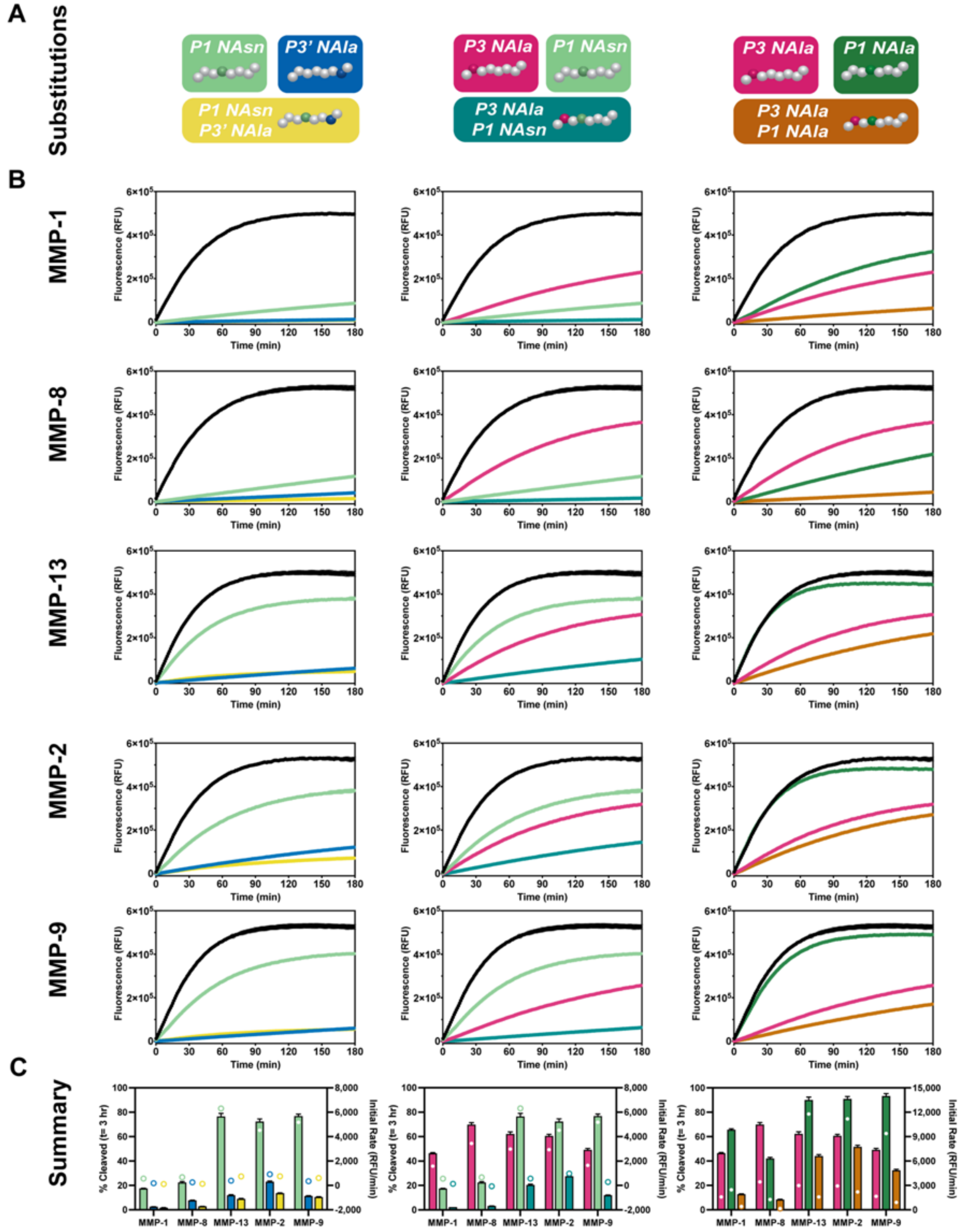

**Figure S7: Tandem substitutions fluorescence traces and summary.** (A) Cartoon illustrations of individual substitutions combined to into three tandem substituted substrates. (B) Fluorescence curves measuring cleavage of each substrate over the course of three hours of exposure to MMPs. (C) Comparison of percent hydrolysis after three hours (bars, left axes) and initial rate of cleavage (unfilled circles, right axes).

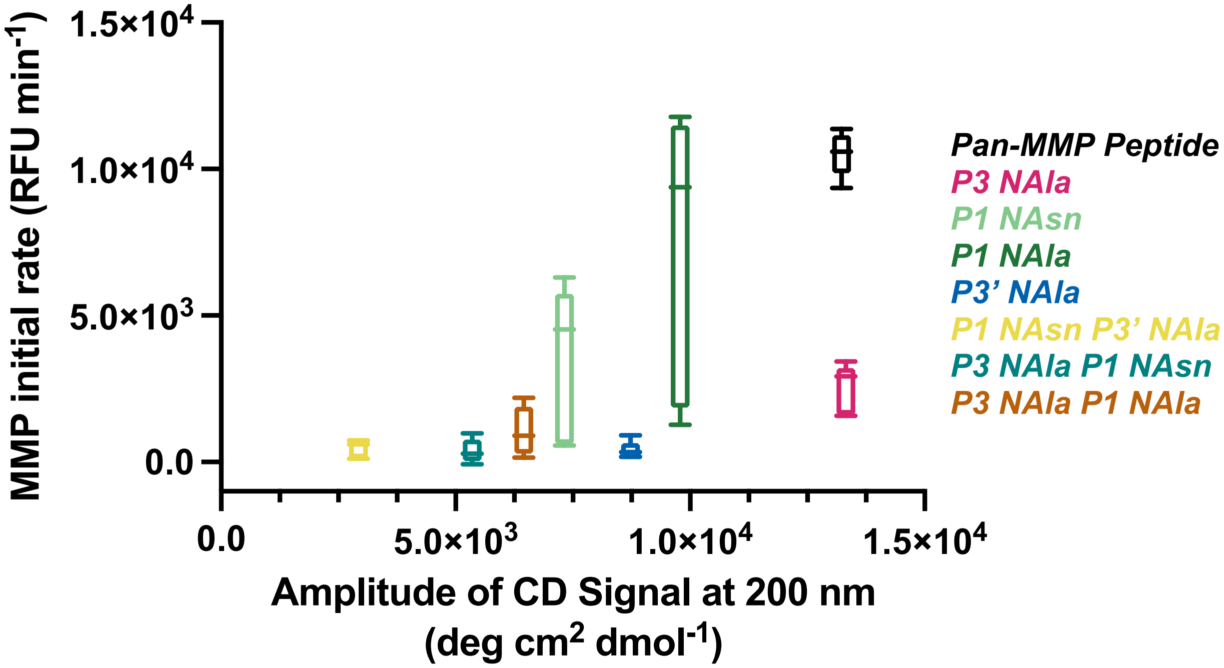

**<u>Figure S8</u>: Correlation between initial rates and secondary structure of tandem substrate library.** Amplitude of the CD peak is correlated with the initial rates of cleavage by the five MMPs.

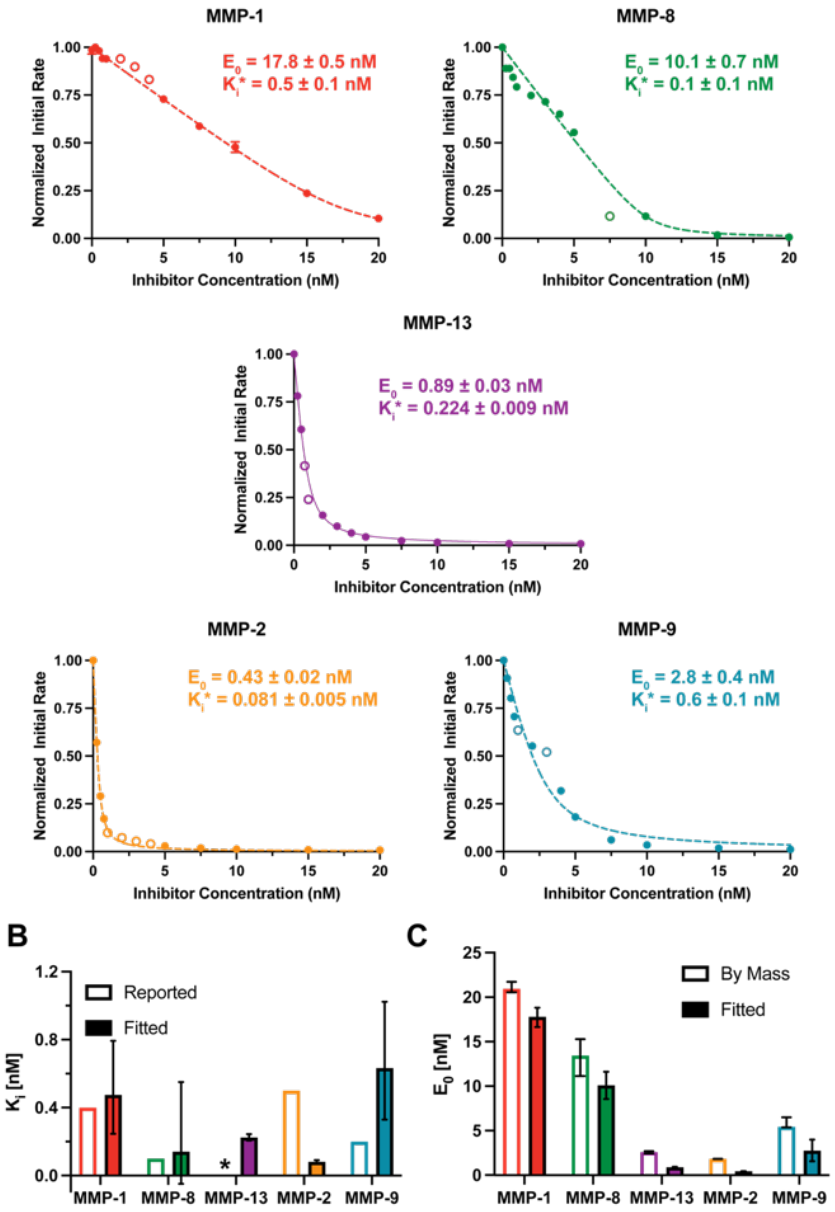

**<u>Figure S9</u>: Active site titrations.** (A) MMPs were titrated using GM 6001 to determine concentration of active enzyme for kinetic parameter fitting. MMPs were incubated with a range of inhibitor concentrations, then the initial rate of cleavage for the *Pan-MMP Peptide* was measured and normalized by the uninhibited velocity. Curves were modeled by the Morison equation for tight binding inhibition with the active enzyme concentration ($E_0$) and applied dissociation constant ($K_i$*) as fitting parameters. Unfilled datapoints were determined to be outliers and were excluded from curve fitting. (B) Comparison of reported $K_i$ concentrations of GM 6001 with MMPs versus the applied $K_i$* determined with the substrate present. *MMP-13 does not have a reported $K_i$ for GM 6001. (C) Expected enzyme concentration calculated from the mass concentration and predicted molecular weight range, versus the concentrations determined by active site titration.

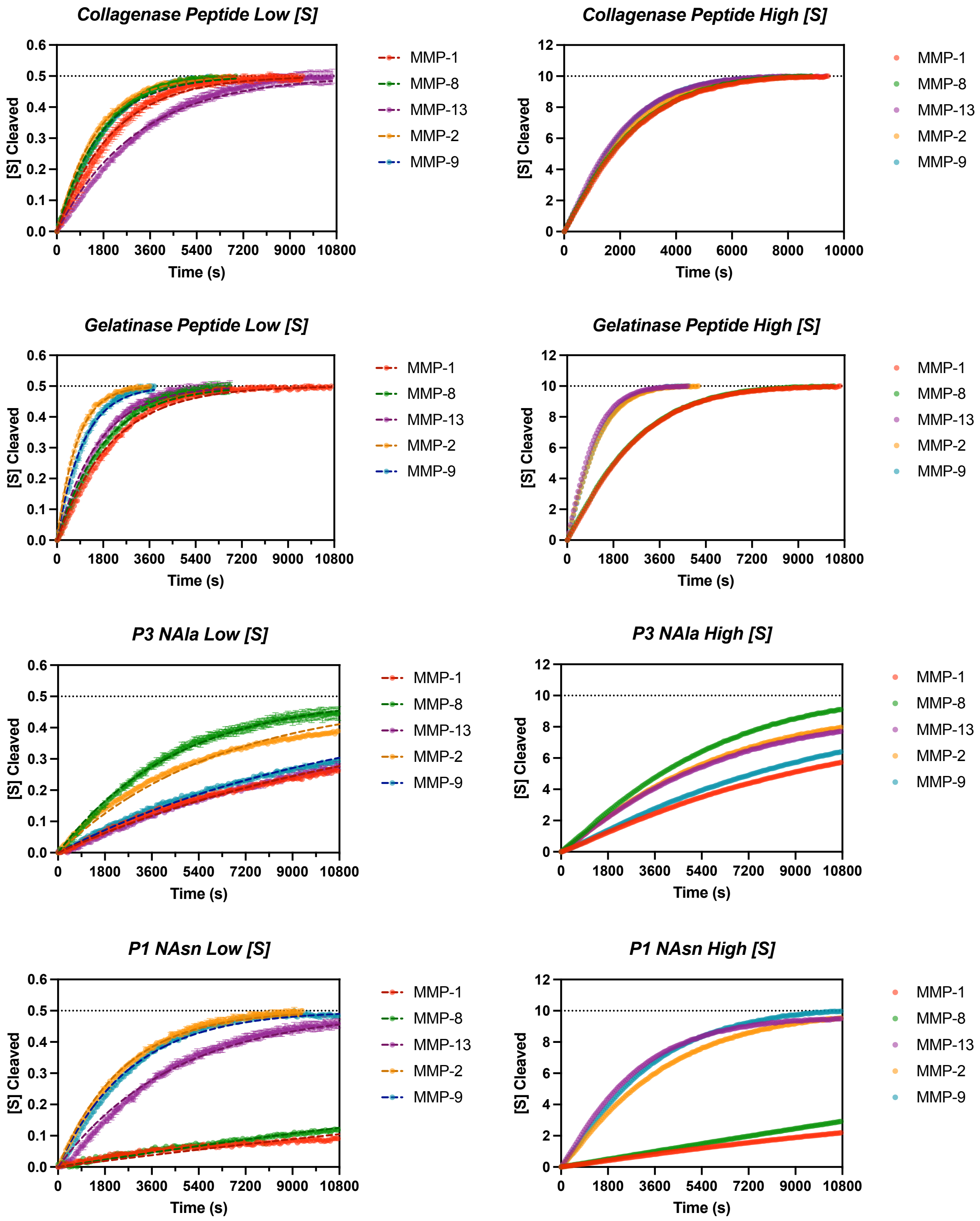
Collagenase Peptide Low [S]
Collagenase Peptide High [S]
Gelatinase Peptide Low [S]
Gelatinase Peptide High [S]
P3 NAla Low [S]
P3 NAla High [S]
P1 NAsn Low [S]
P1 NAsn High [S]
[S] Cleaved
Time (s)
0.0
0.1
0.2
0.3
0.4
0.5
0.6
0
2
4
6
8
10
12
0
1800
3600
5400
7200
9000
10800
0
2000
4000
6000
8000
10000
MMP-1
MMP-8
MMP-13
MMP-2
MMP-9

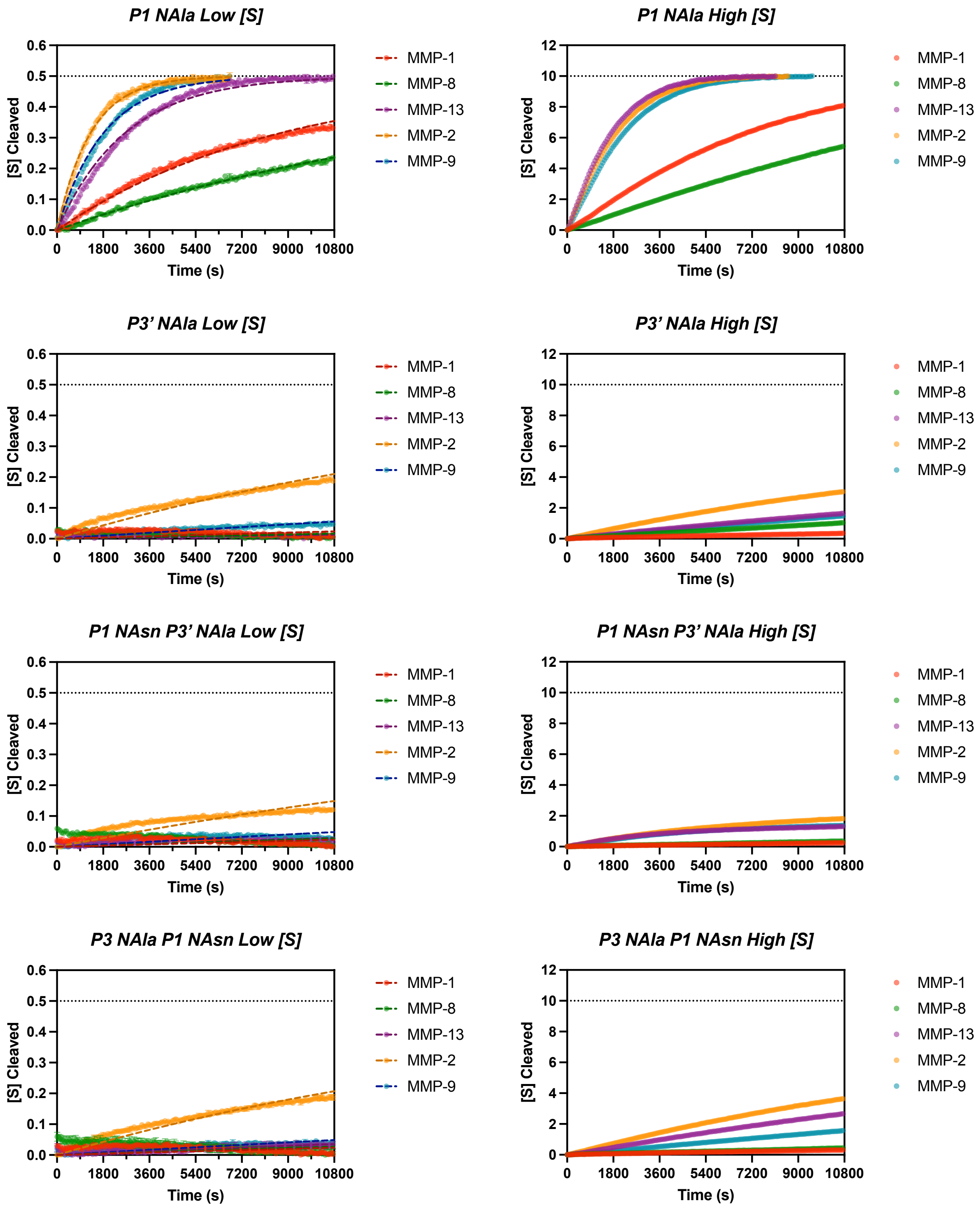
P1 NAla Low [S]
P1 NAla High [S]
P3' NAla Low [S]
P3' NAla High [S]
P1 NAsn P3' NAla Low [S]
P1 NAsn P3' NAla High [S]
P3 NAla P1 NAsn Low [S]
P3 NAla P1 NAsn High [S]
[S] Cleaved
Time (s)
0.0
0.1
0.2
0.3
0.4
0.5
0.6
0
2
4
6
8
10
12
0
1800
3600
5400
7200
9000
10800
MMP-1
MMP-8
MMP-13
MMP-2
MMP-9

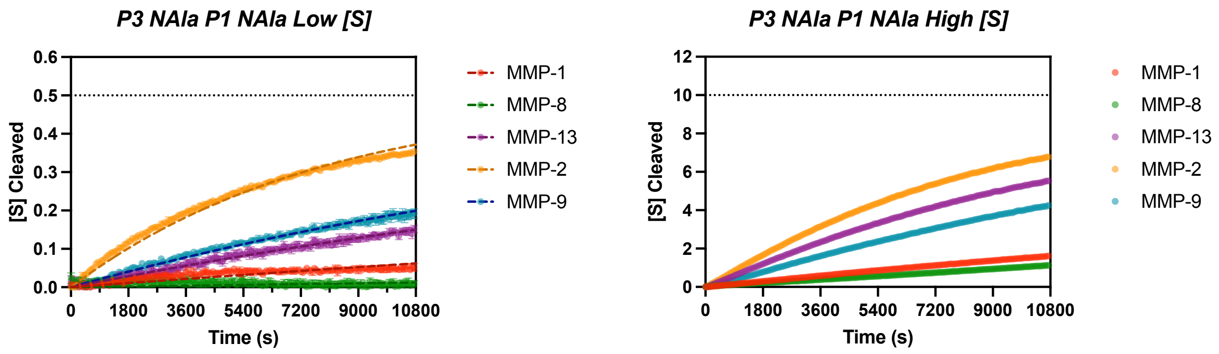

**Figure S10: Substrate degradation screenings for kinetic parameter determination.** The peptide and peptomer substrates comprising the tandem substitution library were screened at low (0.5 µM, left) and high (10 µM, right) substrate concentrations. Relative fluorescence units were converted to product concentration using the fluorescence value corresponding to complete cleavage. The low [S] traces were fit according to the low substrate concentration approximation in order to determine the first-order rate constant ($k_{obs}$). This rate constant was then divided by the enzyme concentration determined by active site titrations to produce an estimated catalytic efficiency. Both the low and high substrate concentration progress curves were used for determining kinetic constants with the tQSSM using the R program available at https://cran.r-project.org/web/packages/EKMCMC/.[2]

**<u>Table S2</u>: Kinetic parameters calculated using the tQSSM.** Substrate traces at 0.5 $\mu$M and 10 $\mu$M, along with substrate and enzyme concentration (as determined by active site titration) were input in R program. Values reported are the posterior mean and standard deviation with default parameters maintained. Samples for which the code did not converge are labeled as not determined (ND).

| Substrate | MMP | tQSSM Approximation | | | | | |
|---|---|---|---|---|---|---|---|
| | | $K_M$ ($\mu$M) | $K_M$ SD | $k_{cat}$ ($s^{-1}$) | $k_{cat}$ SD | Catalytic Efficiency ($M^{-1}$ $s^{-1}$) | Catalytic Efficiency SD |
| *Pan-MMP Peptide* | MMP-1 | 13 | ± 2 | 0.46 | ± 0.05 | 34,000 | ± 6,000 |
| | MMP-8 | 11 | ± 1 | 0.72 | ± 0.07 | 70,000 | ± 10,000 |
| | MMP-13 | 23 | ± 4 | 16 | ± 3 | 700,000 | ± 200,000 |
| | MMP-2 | 10 | ± 2 | 17 | ± 2 | 1,700,000 | ± 300,000 |
| | MMP-9 | 11 | ± 1 | 2.9 | ± 0.3 | 270,000 | ± 40,000 |
| *Gelatinase Peptide* | MMP-1 | 13 | ± 2 | 0.41 | ± 0.04 | 32,000 | ± 5,000 |
| | MMP-8 | 12 | ± 2 | 0.7 | ± 0.07 | 60,000 | ± 10,000 |
| | MMP-13 | 21 | ± 6 | 60 | ± 20 | 3,000,000 | ± 1,000,000 |
| | MMP-2 | 21 | ± 6 | 60 | ± 10 | 2,600,000 | ± 900,000 |
| | MMP-9 | 13 | ± 2 | 5.9 | ± 0.8 | 500,000 | ± 100,000 |
| *P3 NAla* | MMP-1 | 300 | ± 200 | 1.5 | ± 0.9 | 5,000 | ± 4,000 |
| | MMP-8 | 13 | ± 2 | 0.37 | ± 0.04 | 28,000 | ± 6,000 |
| | MMP-13 | 400 | ± 200 | 60 | ± 30 | 150,000 | ± 90,000 |
| | MMP-2 | 130 | ± 30 | 50 | ± 10 | 400,000 | ± 100,000 |
| | MMP-9 | 80 | ± 30 | 3 | ± 1 | 40,000 | ± 20,000 |
| *P1 NAsn* | MMP-1 | 270 | ± 80 | 0.4 | ± 0.1 | 1,300 | ± 600 |
| | MMP-8 | 200 | ± 100 | 0.5 | ± 0.4 | 3,000 | ± 3,000 |
| | MMP-13 | 300 | ± 200 | 120 | ± 50 | 400,000 | ± 200,000 |
| | MMP-2 | 18 | ± 3 | 14 | ± 2 | 800,000 | ± 200,000 |
| | MMP-9 | 10 | ± 1 | 1.7 | ± 0.1 | 180,000 | ± 30,000 |
| *P1 NAla* | MMP-1 | 28 | ± 7 | 0.27 | ± 0.06 | 10,000 | ± 3,000 |
| | MMP-8 | 40 | ± 10 | 0.31 | ± 0.09 | 8,000 | ± 4,000 |
| | MMP-13 | 31 | ± 10 | 24 | ± 7 | 800,000 | ± 30,0000 |
| | MMP-2 | 12 | ± 2 | 22 | ± 3 | 1,800,000 | ± 400,000 |
| | MMP-9 | 12 | ± 2 | 2.9 | ± 0.3 | 240,000 | ± 50,000 |
| *P3' NAla* | MMP-1 | 5 | ± 2 | 0.0028 | ± 0.0005 | 500 | ± 200 |
| | MMP-8 | 140 | ± 90 | 0.15 | ± 0.09 | 1,100 | ± 900 |
| | MMP-13 | 60 | ± 30 | 1.3 | ± 0.6 | 20,000 | ± 10,000 |
| | MMP-2 | 50 | ± 20 | 4 | ± 1 | 90,000 | ± 50,000 |
| | MMP-9 | 300 | ± 200 | 1.5 | ± 0.8 | 5,000 | ± 4,000 |
| *P1 NAsn P3' NAla* | MMP-1 | ND | | | | | |
| | MMP-8 | ND | | | | | |
| | MMP-13 | 1100 | ± 500 | 16 | ± 8 | 10,000 | ± 10,000 |
| | MMP-2 | 200 | ± 100 | 8 | ± 5 | 50,000 | ± 40,000 |
| | MMP-9 | 160 | ± 40 | 0.9 | ± 0.2 | 5,000 | ± 2,000 |
| *P3 NAla P1 NAsn* | MMP-1 | 160 | ± 80 | 0.03 | ± 0.01 | 200 | ± 100 |
| | MMP-8 | ND | | | | | |
| | MMP-13 | 500 | ± 200 | 16 | ± 5 | 30,000 | ± 10,000 |
| | MMP-2 | 110 | ± 30 | 12 | ± 3 | 110,000 | ± 40,000 |
| | MMP-9 | 2000 | ± 2000 | 10 | ± 10 | 6,000 | ± 7,000 |
| *P3 NAla P1 NAla* | MMP-1 | 300 | ± 200 | 0.3 | ± 0.1 | 900 | ± 700 |
| | MMP-8 | ND | | | | | |
| | MMP-13 | 900 | ± 500 | 70 | ± 40 | 80,000 | ± 70,000 |
| | MMP-2 | 200 | ± 100 | 60 | ± 30 | 300,000 | ± 200,000 |
| | MMP-9 | 130 | ± 50 | 2.4 | ± 1 | 20,000 | ± 10,000 |

**<u>Table S3</u>: Kinetic parameters calculated using the low substrate concentration approximation.** Catalytic efficiency is determined using the $k_{obs}$ exponential fitting parameter from the 0.5 $\mu$M traces in **Figure S10**. Values reported are from fits from three averaged replicates and include the parameter mean and standard error.

| Substrate | MMP | Low Substrate Concentration Approximation | | | |
|---|---|---|---|---|---|
| | | $k_{obs}$ ($s^{-1}$) | $k_{obs}$ SE | Catalytic Efficiency ($M^{-1}$ $s^{-1}$) | Catalytic Efficiency SE |
| *Pan-MMP Peptide* | MMP-1 | 4.69E-4 | 4E-6 | 26,300 | ± 800 |
| | MMP-8 | 5.96E-4 | 5E-6 | 59,000 | ± 4,000 |
| | MMP-13 | 3.23E-4 | 3E-6 | 363,000 | ± 5,000 |
| | MMP-2 | 6.63E-4 | 3E-6 | 1,540,000 | ± 70,000 |
| | MMP-9 | 5.89E-4 | 6E-6 | 210,000 | ± 30,000 |
| *Gelatinase Peptide* | MMP-1 | 4.75E-4 | 4E-6 | 26,700 | ± 800 |
| | MMP-8 | 5.45E-4 | 5E-6 | 54,000 | ± 4,000 |
| | MMP-13 | 6.21E-4 | 6E-6 | 700,000 | ± 10,000 |
| | MMP-2 | 1.3E-3 | 1E-5 | 3,000,000 | ± 100,000 |
| | MMP-9 | 9.8E-4 | 1E-5 | 350,000 | ± 50,000 |
| *P3 NAla* | MMP-1 | 7.47E-5 | 4E-7 | 4,200 | ± 100 |
| | MMP-8 | 2.22E-4 | 8E-7 | 22,000 | ± 2,000 |
| | MMP-13 | 7.46E-5 | 2E-7 | 83,800 | ± 900 |
| | MMP-2 | 1.599E-4 | 1E-6 | 370,000 | ± 20,000 |
| | MMP-9 | 8.66E-5 | 3E-7 | 31,000 | ± 4,000 |
| *P1 NAsn* | MMP-1 | 2.18E-5 | 3E-7 | 1,220 | ± 40 |
| | MMP-8 | 2.68E-5 | 2E-7 | 2,600 | ± 200 |
| | MMP-13 | 2.25E-4 | 1E-6 | 253,000 | ± 3,000 |
| | MMP-2 | 4.03E-4 | 2E-6 | 940,000 | ± 40,000 |
| | MMP-9 | 3.61E-4 | 2E-6 | 130,000 | ± 20,000 |
| *P1 NAla* | MMP-1 | 1.142E-4 | 5E-7 | 6,400 | ± 200 |
| | MMP-8 | 5.92E-5 | 2E-7 | 5,900 | ± 400 |
| | MMP-13 | 3.76E-4 | 3E-6 | 422,000 | ± 6,000 |
| | MMP-2 | 7.62E-4 | 5E-6 | 1,770,000 | ± 80,000 |
| | MMP-9 | 5.5E-4 | 5E-6 | 200,000 | ± 30,000 |
| *P3' NAla* | MMP-1 | 4.5E-6 | 4E-7 | 250 | ± 20 |
| | MMP-8 | 3.1E-6 | 3E-7 | 310 | ± 40 |
| | MMP-13 | 2.4E-6 | 2E-7 | 2,800 | ± 200 |
| | MMP-2 | 5.04E-5 | 5E-7 | 117,000 | ± 6,000 |
| | MMP-9 | 1.09E-5 | 2E-7 | 3,900 | ± 600 |
| *P1 NAsn P3' NAla* | MMP-1 | 4.5E-6 | 4E-7 | 250 | ± 20 |
| | MMP-8 | 5.2E-6 | 6E-7 | 520 | ± 70 |
| | MMP-13 | 5.6E-6 | 2E-7 | 6,300 | ± 200 |
| | MMP-2 | 3.28E-5 | 6E-7 | 76,000 | ± 4,000 |
| | MMP-9 | 9.4E-6 | 3E-7 | 3,300 | ± 500 |
| *P3 NAla P1 NAsn* | MMP-1 | 4.5E-6 | 4E-7 | 250 | ± 30 |
| | MMP-8 | 5.2E-6 | 6E-7 | 520 | ± 70 |
| | MMP-13 | 6.8E-6 | 1E-7 | 7,700 | ± 200 |
| | MMP-2 | 4.95E-5 | 4E-7 | 115,000 | ± 5,000 |
| | MMP-9 | 9.5E-6 | 2E-7 | 3,400 | ± 500 |
| *P3 NAla P1 NAla* | MMP-1 | 1.23E-5 | 2E-7 | 690 | ± 20 |
| | MMP-8 | 2.3E-6 | 2E-7 | 230 | ± 20 |
| | MMP-13 | 3.3E-5 | 2E-7 | 37,100 | ± 400 |
| | MMP-2 | 1.261E-4 | 7E-7 | 290,000 | ± 10,000 |
| | MMP-9 | 4.71E-5 | 2E-7 | 17,000 | ± 2000 |

**<u>Table S4</u>: Catalytic efficiency comparison.** Comparison of the catalytic efficiencies approximated by the two methods shows good agreement. Catalytic efficiencies are also benchmarked versus the *Pan-MMP Peptide* to demonstrate the inherent variability in activity of the MMPs and the effect that peptoid substitutions have on catalytic efficiency.

| Substrate | MMP | tQSSM Approximation | | | Low Substrate Concentration Approximation | | |
|---|---|---|---|---|---|---|---|
| | | Catalytic Efficiency ($M^{-1}$ $s^{-1}$) | Catalytic Efficiency SD | % of Pan-MMP Peptide Catalytic Efficiency | Catalytic Efficiency ($M^{-1}$ $s^{-1}$) | Catalytic Efficiency SE | % of Pan-MMP Peptide Catalytic Efficiency |
| *Pan-MMP Peptide* | MMP-1 | 34,000 | ± 6,000 | - | 26,300 | ± 800 | - |
| | MMP-8 | 70,000 | ± 10,000 | - | 59,000 | ± 4,000 | - |
| | MMP-13 | 700,000 | ± 200,000 | - | 363,000 | ± 5,000 | - |
| | MMP-2 | 1,700,000 | ± 300,000 | - | 1,540,000 | ± 70,000 | - |
| | MMP-9 | 270,000 | ± 40,000 | - | 210,000 | ± 30,000 | - |
| *Gelatinase Peptide* | MMP-1 | 32,000 | ± 5,000 | 94% | 26,700 | ± 800 | 102% |
| | MMP-8 | 60,000 | ± 10,000 | 86% | 54,000 | ± 4,000 | 92% |
| | MMP-13 | 3,000,000 | ± 1,000,000 | 429% | 700,000 | ± 10,000 | 193% |
| | MMP-2 | 2,600,000 | ± 900,000 | 153% | 3,000,000 | ± 100,000 | 195% |
| | MMP-9 | 500,000 | ± 100,000 | 185% | 350,000 | ± 50,000 | 167% |
| *P3 NAla* | MMP-1 | 5,000 | ± 4,000 | 15% | 4,200 | ± 100 | 15% |
| | MMP-8 | 28,000 | ± 6,000 | 40% | 22,000 | ± 2,000 | 40% |
| | MMP-13 | 150,000 | ± 90,000 | 21% | 83,800 | ± 900 | 21% |
| | MMP-2 | 400,000 | ± 100,000 | 24% | 370,000 | ± 20,000 | 24% |
| | MMP-9 | 40,000 | ± 20,000 | 15% | 31,000 | ± 4,000 | 15% |
| *P1 NAsn* | MMP-1 | 1,300 | ± 600 | 4% | 1,220 | ± 40 | 15% |
| | MMP-8 | 3,000 | ± 3,000 | 4% | 2,600 | ± 200 | 40% |
| | MMP-13 | 400,000 | ± 200,000 | 57% | 253,000 | ± 3,000 | 21% |
| | MMP-2 | 800,000 | ± 200,000 | 47% | 940,000 | ± 40,000 | 24% |
| | MMP-9 | 180,000 | ± 30,000 | 67% | 130,000 | ± 20,000 | 15% |
| *P1 NAla* | MMP-1 | 10,000 | ± 3,000 | 29% | 6,400 | ± 200 | 15% |
| | MMP-8 | 8,000 | ± 4,000 | 11% | 5,900 | ± 400 | 40% |
| | MMP-13 | 800,000 | ± 30,0000 | 114% | 422,000 | ± 6,000 | 21% |
| | MMP-2 | 1,800,000 | ± 400,000 | 106% | 1,770,000 | ± 80,000 | 24% |
| | MMP-9 | 240,000 | ± 50,000 | 89% | 200,000 | ± 30,000 | 15% |
| *P3' NAla* | MMP-1 | 500 | ± 200 | 1% | 250 | ± 20 | 15% |
| | MMP-8 | 1,100 | ± 900 | 2% | 310 | ± 40 | 40% |
| | MMP-13 | 20,000 | ± 10,000 | 3% | 2,800 | ± 200 | 21% |
| | MMP-2 | 90,000 | ± 50,000 | 5% | 117,000 | ± 6,000 | 24% |
| | MMP-9 | 5,000 | ± 4,000 | 2% | 3,900 | ± 600 | 15% |
| *P1 NAsn P3' NAla* | MMP-1 | | ND | | 250 | ± 20 | 15% |
| | MMP-8 | | ND | | 520 | ± 70 | 40% |
| | MMP-13 | 10,000 | ± 10,000 | 1% | 6,300 | ± 200 | 21% |
| | MMP-2 | 50,000 | ± 40,000 | 3% | 76,000 | ± 4,000 | 24% |
| | MMP-9 | 5,000 | ± 2,000 | 2% | 3,300 | ± 500 | 15% |
| *P3 NAla P1 NAsn* | MMP-1 | 200 | ± 100 | 1% | 250 | ± 30 | 15% |
| | MMP-8 | | ND | | 520 | ± 70 | 40% |
| | MMP-13 | 30,000 | ± 10,000 | 4% | 7,700 | ± 200 | 21% |
| | MMP-2 | 110,000 | ± 40,000 | 6% | 115,000 | ± 5,000 | 24% |
| | MMP-9 | 6,000 | ± 7,000 | 2% | 3,400 | ± 500 | 15% |
| *P3 NAla P1 NAla* | MMP-1 | 900 | ± 700 | 3% | 690 | ± 20 | 15% |
| | MMP-8 | | ND | | 230 | ± 20 | 40% |
| | MMP-13 | 80,000 | ± 70,000 | 11% | 37,100 | ± 400 | 21% |
| | MMP-2 | 300,000 | ± 200,000 | 18% | 290,000 | ± 10,000 | 24% |
| | MMP-9 | 20,000 | ± 10,000 | 7% | 17,000 | ± 2000 | 15% |

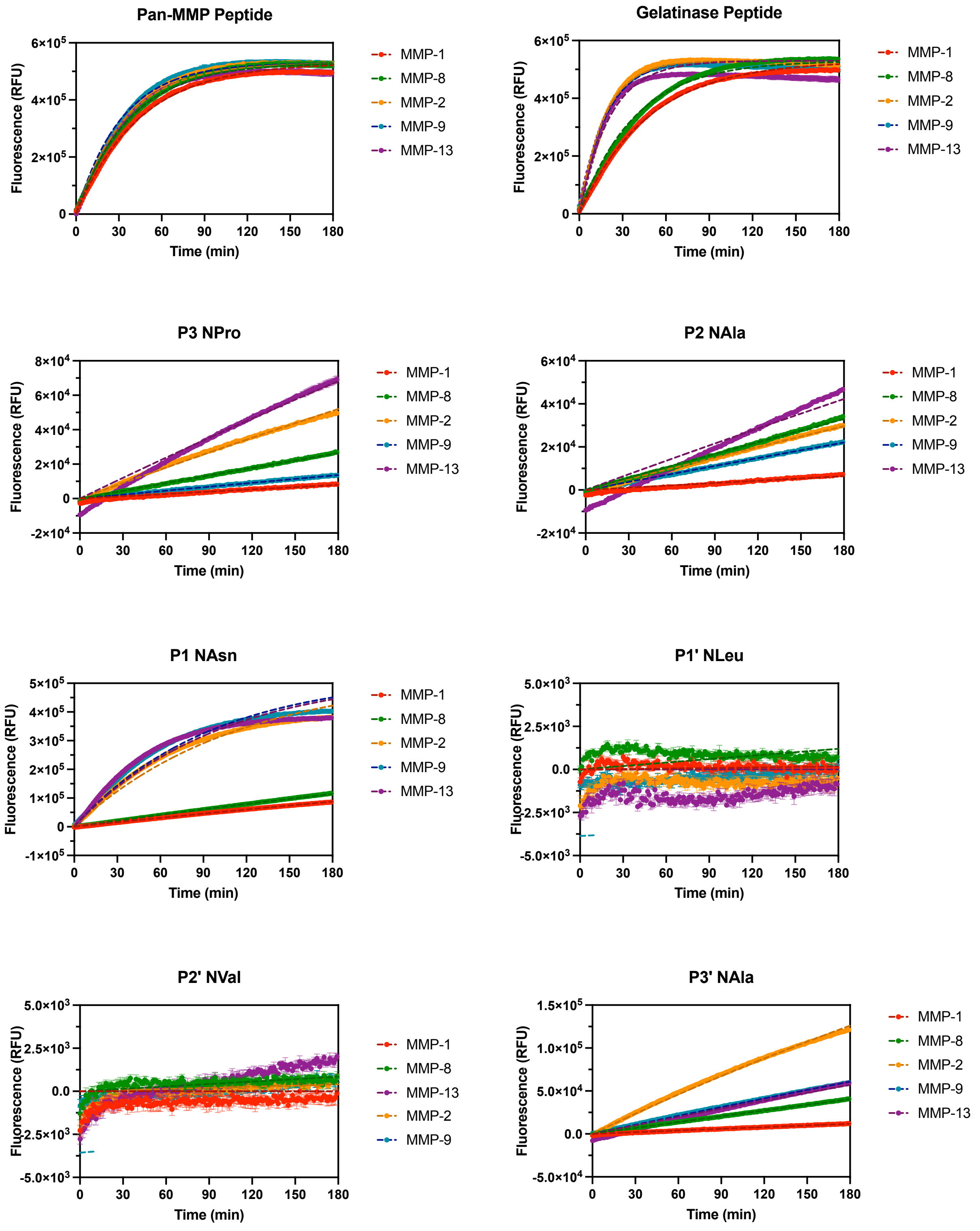

Pan-MMP Peptide
Gelatinase Peptide
P3 NPro
P2 NAla
P1 NAsn
P1' NLeu
P2' NVal
P3' NAla
Fluorescence (RFU)
Time (min)
MMP-1
MMP-8
MMP-2
MMP-9
MMP-13

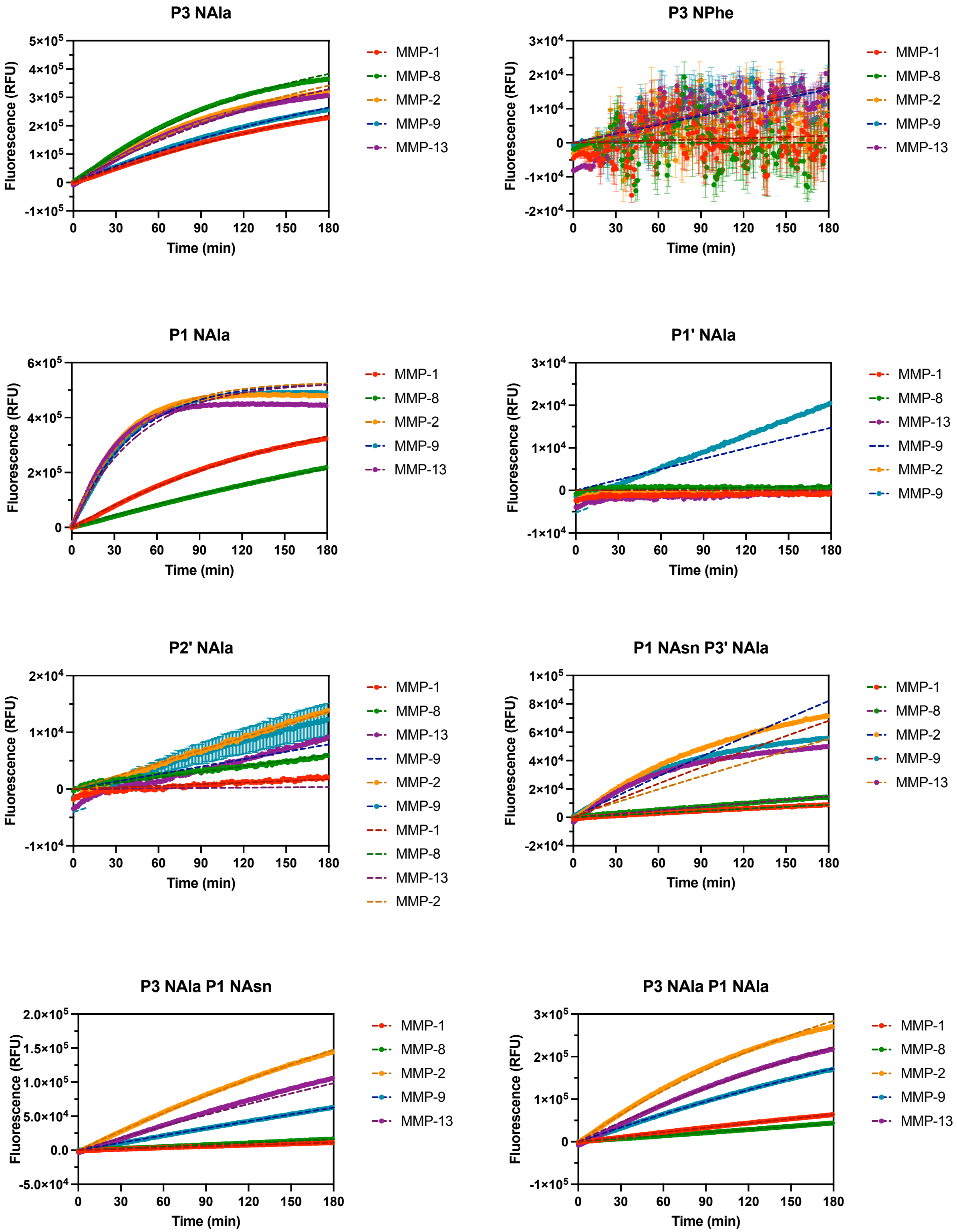

**Figure S11: Exponential plateau function fits.** Fluorescence data (points) and fitted curves (dashed lines) for each substrate by MMP. Fits were constrained to an initial value of 0 and a plateau value of 530,000 RFU.

**<u>Table S5</u>: log(k) values used for multivariate data analysis.**

| | *Collagenase Peptide* | *Gelatinase Peptide* | *P3 NPro* | *P2 NAla* | *P1 NAsn* | *P3' NAla* | *P3 NAla* | *P1 NAla* | *P1 NAsn P3' NAla* | *P3 NAla P1 NAsn* | *P3 NAla P1 NAla* |
|---|---|---|---|---|---|---|---|---|---|---|---|
| **MMP-1** | -1.62 | -1.67 | -4.12 | -4.19 | -3.00 | -3.92 | -2.48 | -2.26 | -4.00 | -3.93 | -3.13 |
| **MMP-1** | -1.66 | -1.68 | -4.05 | -4.14 | -3.00 | -3.92 | -2.48 | -2.27 | -4.03 | -3.92 | -3.15 |
| **MMP-1** | -1.64 | -1.68 | -4.06 | -4.17 | -3.00 | -3.87 | -2.49 | -2.26 | -4.04 | -3.93 | -3.15 |
| **MMP-8** | -1.57 | -1.59 | -3.53 | -3.45 | -2.86 | -3.35 | -2.14 | -2.54 | -3.79 | -3.77 | -3.32 |
| **MMP-8** | -1.60 | -1.58 | -3.55 | -3.45 | -2.89 | -3.37 | -2.15 | -2.54 | -3.81 | -3.76 | -3.34 |
| **MMP-8** | -1.55 | -1.58 | -3.56 | -3.44 | -2.86 | -3.34 | -2.15 | -2.55 | -3.81 | -3.77 | -3.33 |
| **MMP-13** | -1.60 | -1.32 | -3.13 | -3.35 | -2.01 | -3.21 | -2.27 | -1.67 | -3.21 | -2.94 | -2.52 |
| **MMP-13** | -1.58 | -1.36 | -3.12 | -3.34 | -2.01 | -3.20 | -2.28 | -1.69 | -3.25 | -2.95 | -2.53 |
| **MMP-13** | -1.54 | -1.29 | -3.10 | -3.32 | -1.97 | -3.18 | -2.26 | -1.64 | -3.21 | -2.93 | -2.51 |
| **MMP-2** | -1.56 | -1.28 | -3.25 | -3.50 | -2.06 | -2.83 | -2.24 | -1.61 | -3.03 | -2.74 | -2.36 |
| **MMP-2** | -1.53 | -1.22 | -3.23 | -3.50 | -2.07 | -2.83 | -2.24 | -1.62 | -3.03 | -2.75 | -2.38 |
| **MMP-2** | -1.53 | -1.23 | -3.24 | -3.48 | -2.03 | -2.81 | -2.24 | -1.58 | -3.03 | -2.73 | -2.37 |
| **MMP-9** | -1.48 | -1.25 | -3.86 | -3.64 | -1.99 | -3.18 | -2.41 | -1.62 | -3.11 | -3.16 | -2.65 |
| **MMP-9** | -1.54 | -1.30 | -3.83 | -3.61 | -1.97 | -3.17 | -2.43 | -1.63 | -3.13 | -3.15 | -2.66 |
| **MMP-9** | -1.48 | -1.24 | -3.82 | -3.62 | -1.97 | -3.16 | -2.41 | -1.65 | -3.11 | -3.16 | -2.67 |

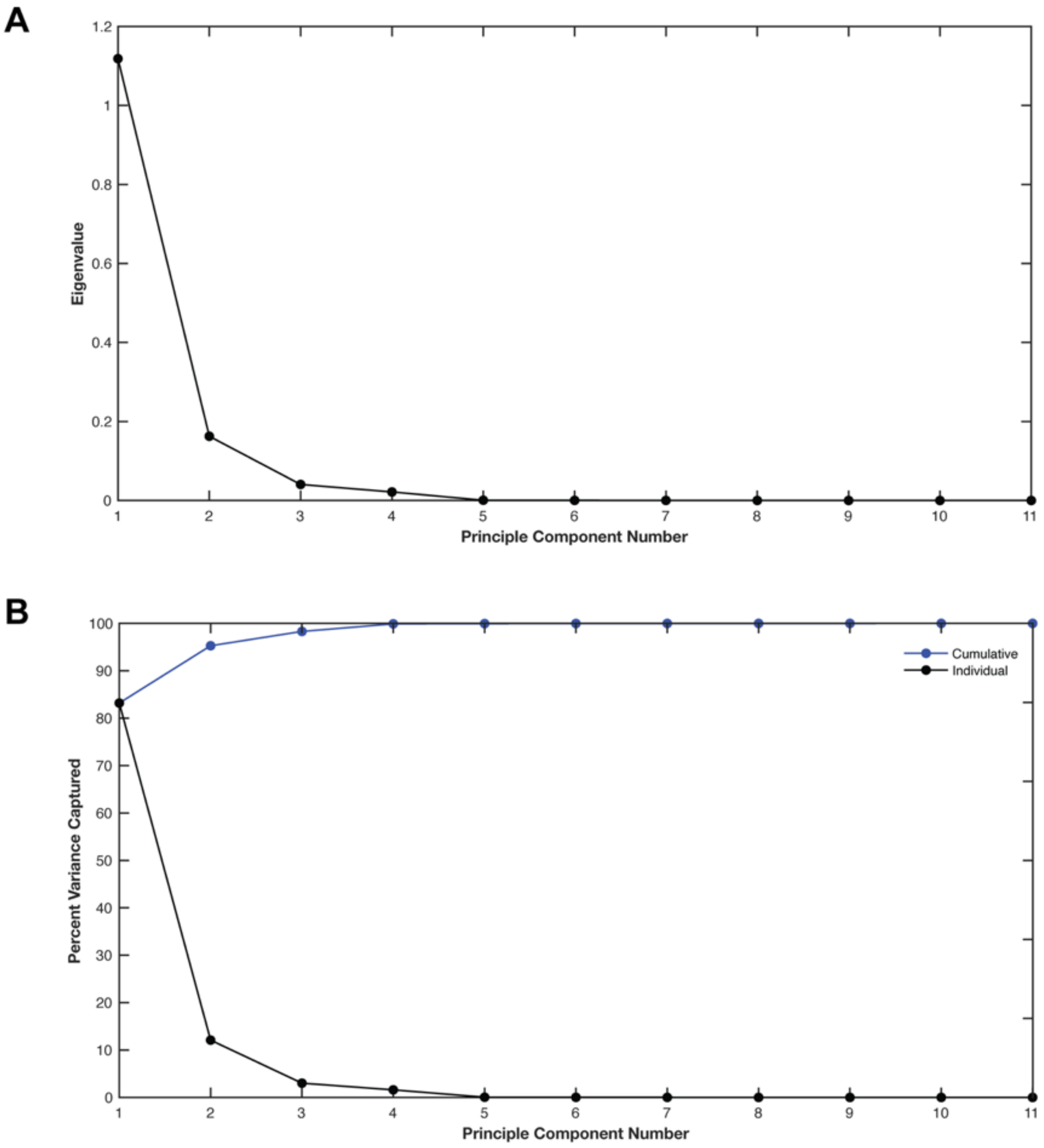

**<u>Figure S12</u>: Scree plots.** Variance explained by each principal component in terms of (A) eigenvalues and (B) percentages.

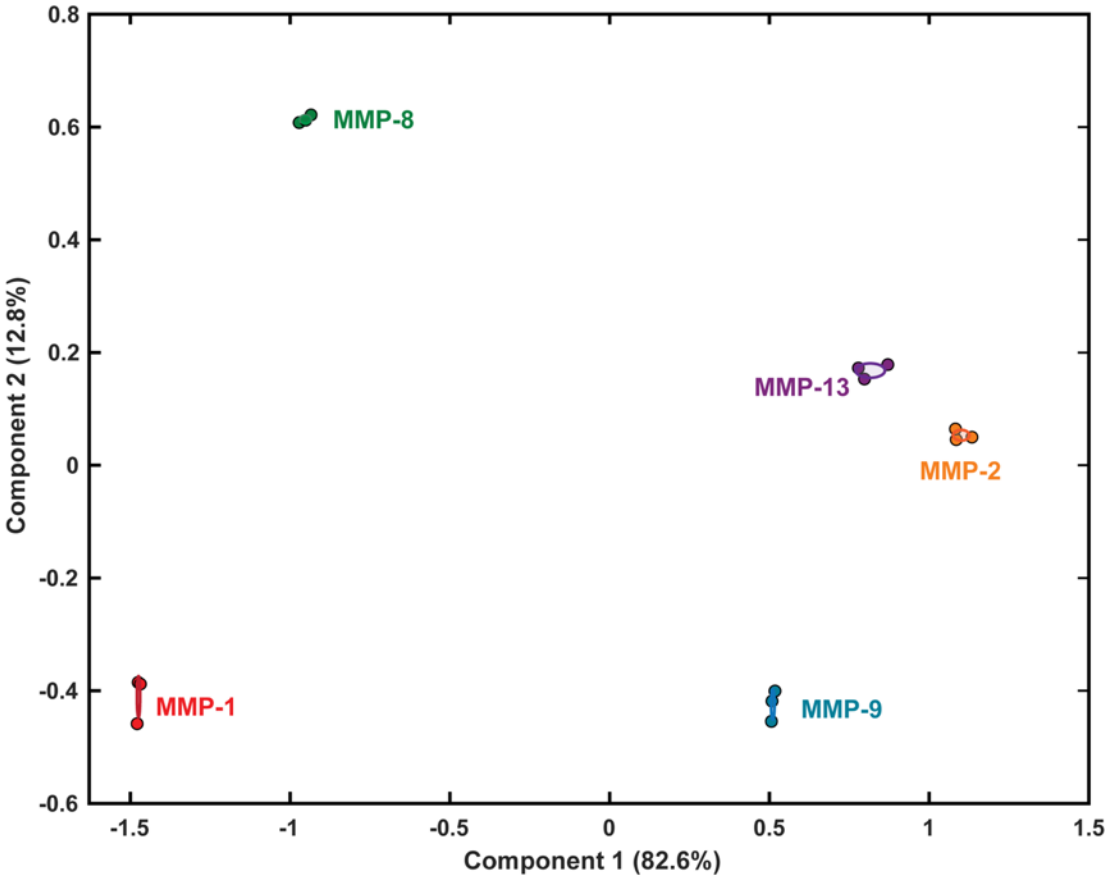

**<u>Figure S13</u>: Two dimensional PCA plot with 95% confidence intervals.**

**Table S6: Data reconstructed by PCA.**

| | Principal components | | | Reconstructed Data | | |
|---|---|---|---|---|---|---|
| | PC1 | PC2 | PC3 | C1 | C2 | C3 |
| MMP-1 | -1.479 | -0.459 | 0.149 | -1.62 | -1.67 | -4.12 |
| MMP-1 | -1.475 | -0.385 | 0.186 | -1.66 | -1.68 | -4.05 |
| MMP-1 | -1.469 | -0.388 | 0.167 | -1.64 | -1.681 | -4.06 |
| MMP-8 | -0.934 | 0.622 | -0.153 | -1.57 | -1.59 | -3.53 |
| MMP-8 | -0.971 | 0.608 | -0.139 | -1.60 | -1.58 | -3.55 |
| MMP-8 | -0.951 | 0.612 | -0.176 | -1.55 | -1.58 | -3.56 |
| MMP-13 | 0.797 | 0.153 | 0.216 | -1.60 | -1.32 | -3.13 |
| MMP-13 | 0.779 | 0.172 | 0.222 | -1.58 | -1.36 | -3.12 |
| MMP-13 | 0.871 | 0.178 | 0.201 | -1.54 | -1.29 | -3.10 |
| MMP-2 | 1.085 | 0.045 | 0.079 | -1.56 | -1.28 | -3.25 |
| MMP-2 | 1.082 | 0.064 | 0.070 | -1.53 | -1.22 | -3.23 |
| MMP-2 | 1.134 | 0.050 | 0.057 | -1.53 | -1.23 | -3.24 |
| MMP-9 | 0.506 | -0.454 | -0.299 | -1.48 | -1.25 | -3.86 |
| MMP-9 | 0.508 | -0.418 | -0.274 | -1.54 | -1.30 | -3.83 |
| MMP-9 | 0.517 | -0.400 | -0.306 | -1.48 | -1.24 | -3.82 |

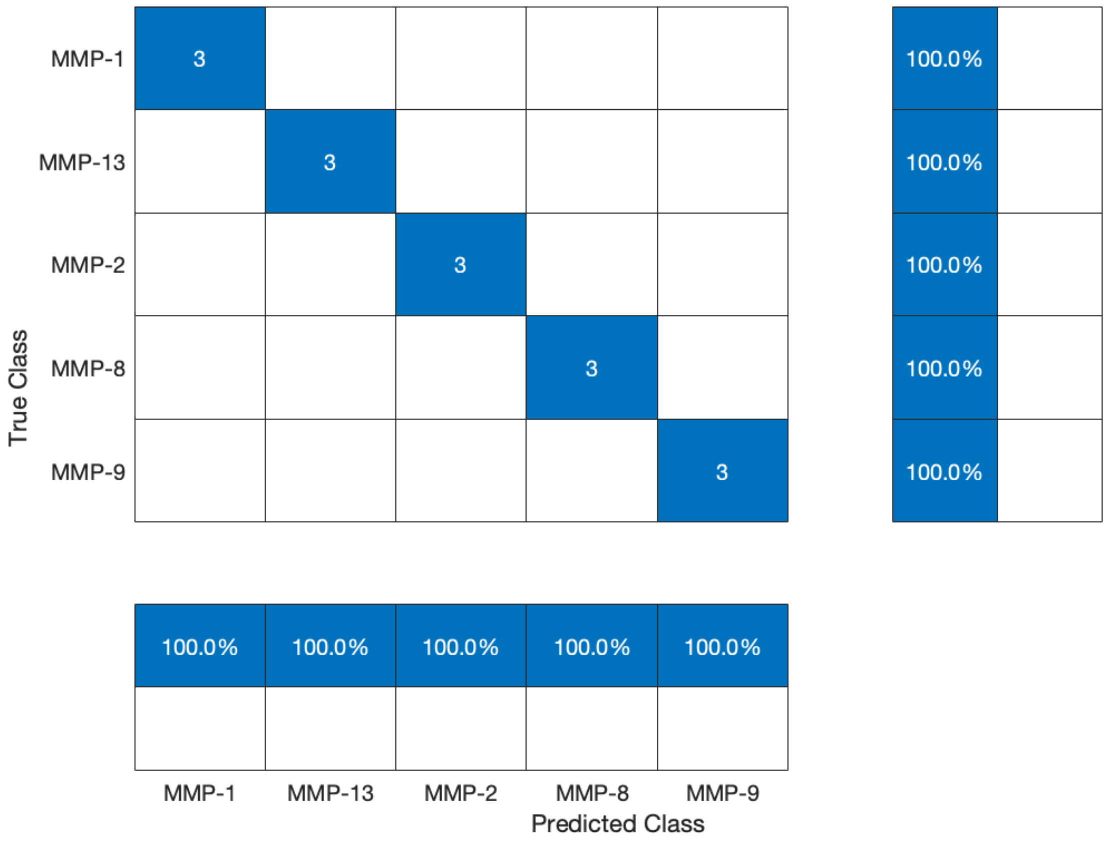

**<u>Figure S14</u>: kNN classification results.** With stratified three-fold cross-validation the kNN model was able to accurately classify all datapoints. The results were the same when the data input was reconstructed variables from PCA, as well as the raw log(k) values.
